# CryoForge: A Self-Correcting Agent for Cryo-EM Model Building That Learns When to Act and When to Stop

**DOI:** 10.64898/2026.08.15.745007

**Authors:** Wei Feng, Yideng Jiang, Fei Sun, Jianyi Yang, Xin Gao, Fa Zhang, Renmin Han

## Abstract

Automated atomic model building has accelerated cryo-EM structure determination, but different builders leave distinct residual error profiles requiring expert inspection. The post-building challenge is to decide which local interpretations are sufficiently supported by experimental evidence to be retained, corrected or rejected. Here we introduce CryoForge, an evidence-gated post-builder agent that separates repair proposal from repair acceptance. Rule- and learning-based components identify candidate regions and prioritize legal actions, whereas an independent evidence gate evaluates each edit using map and half-map support, stereochemistry, connectivity and local structural context. Supported edits are retained; unsupported or conflicting modifications are rejected, rolled back, stopped or escalated for expert review. Across a resolution-stratified benchmark, 84.9% of 26,153 released trajectories yielded standard validated improvements and 3.7% yielded low-confidence partial improvements, with no quality-degrading edit retained in the final promoted models. Relative to rule-only control, learned prioritization reduced non-improving candidates and harmful actions while preserving global structural stability. External evaluations using an alternative initializer, same-team automated/manual-assisted challenge submissions and three recently released complex assemblies showed that CryoForge adapts to distinct residual error phenotypes and performs bounded, evidence-supported correction without uncontrolled remodeling. CryoForge provides a builder-independent, scalable and auditable correction layer between automated model generation and expert structural interpretation.

## Introduction

Single-particle cryo-electron microscopy (cryo-EM) has transformed structural biology by enabling near-atomic-resolution determination of increasingly complex macro-molecular assemblies. Advances in electron detectors, microscope instrumentation, image-processing algorithms and three-dimensional reconstruction have established cryo-EM as a major platform for studying membrane proteins, molecular machines and large macromolecular complexes [1–3]. The biological value of a reconstruction, however, depends not only on the quality of the density map but also on the reliability of the atomic model used to interpret it. Atomic coordinates are not direct experimental observations; they are structural interpretations obtained by combining density information with sequence constraints, chemical principles and prior structural knowledge [4, 5]. Initial model generation is therefore not the endpoint of cryo-EM structure determination, but the beginning of a process of model validation, correction and evidence-based review.

A fundamental difficulty in cryo-EM model interpretation arises from the spatial heterogeneity of the experimental information. Local resolution and interpretability can vary substantially within a single reconstruction [6, 7]: rigid core regions may provide strong constraints for backbone tracing and side-chain placement, whereas flexible loops, terminal segments, molecular interfaces and ligand-binding regions can exhibit weak, incomplete or ambiguous density. Different regions within the same atomic model can consequently have markedly different levels of experimental support, as reflected by local measures of atomic resolvability and map–model agreement [8, 9]. This spatial heterogeneity creates unavoidable residual modeling burden after initial model generation. Some regions may remain missing, disconnected or incompletely interpreted because the experimental information is insufficient, whereas others may contain locally plausible but inaccurate coordinate assignments or interpretations that lack adequate experimental support. Such problems commonly arise where density interpretation, sequence assignment, geometric constraints and local structural context are difficult to reconcile. The central challenge after automated model building is therefore not merely to improve global model scores, but to identify which local interpretations remain unreliable and determine whether and how they should be corrected.

Automated model-building methods have substantially reduced the effort required to convert cryo-EM maps into initial atomic coordinates. Density-driven procedures established automated backbone tracing and sequence assignment, but their performance remains strongly dependent on local map quality and can be limited by incomplete tracing, broken connectivity and missing side-chain interpretation in poorly resolved regions [10]. Learning-based and prior-assisted systems further integrate cryo-EM density with sequence information, learned structural representations and predicted structural priors to improve model completeness, continuity and geometric plausibility [11–13]. CryoAtom, E3-CryoFold and DiffModeler further illustrate the rapid development of end-to-end and prior-assisted modeling strategies [14–16]. Nevertheless, no initial model-generation strategy completely eliminates residual modeling burden. Because different initializers place different emphasis on density evidence, sequence information and structural priors, they leave method-dependent residual modeling phenotypes [17]. Density-driven approaches may remain conservative in weak-density regions and leave missing or disconnected interpretations, whereas prior-assisted approaches can produce more complete models whose local assignments must still be validated against the specific experimental state represented by the target map [18– 20]. The emerging need is therefore not another model builder, but a post-builder correction layer capable of handling residual problems across different initial-modeling paradigms under a common local evidence framework.

Current cryo-EM workflows provide powerful but functionally specialized components. Coot and ISOLDE support interactive inspection and rebuilding [21, 22]; PHENIX and Servalcat optimize existing coordinates against experimental density and structural restraints [23–25]; and MolProbity and EMRinger evaluate stereochemistry and side-chain placement [26, 27], whereas Q-score and FSC-Q assess local map–model support [8, 9]. Recent computational efforts have also automated individual aspects of post-modeling analysis, including residue-level error detection and density-constrained refinement [28, 29]. These components primarily perform distinct functions in model construction, coordinate optimization or quality assessment and do not jointly resolve a central decision problem: which local structural interpretations should be modified, preserved or rejected after initial model generation? Addressing this problem requires a builder-independent post-builder correction layer that identifies regions requiring intervention, selects legal and appropriate repair actions and uses independent evidence to determine whether each modification should enter the final model. Its objective is not to maximize the number of edits, but to preserve reliable structural regions while selectively modifying only local interpretations supported by sufficient experimental and structural evidence.

Here we introduce CryoForge, an evidence-gated post-builder agent for correcting residual problems in cryo-EM initial atomic models. Rather than replacing existing model-generation methods, CryoForge operates after initial model construction across different builders and separates repair proposal from repair acceptance. Rule- and learning-based components identify candidate regions, a GNN/RL policy prioritizes legal repair actions and an independent, predefined and auditable evidence gate determines whether an edit is sufficiently supported to enter the final model. Each modification is evaluated using complementary evidence from the full map and half maps, stereochemical quality, sequence consistency, chain connectivity and local structural context. Supported edits are retained, whereas unsupported or conflicting modifications are rejected, rolled back, stopped or escalated for expert review. A supervisory LLM summarizes evidence and assists with complex cases but does not generate coordinate edits or override failed quality control. By establishing a builder-independent, verifiable and rollback-enabled correction layer between automated model generation and expert structural interpretation, CryoForge provides a unified, scalable and auditable framework for addressing residual local problems left by different initialization workflows.

## Results

### CryoForge overview: evidence-gated local repair of cryo-EM initial models

CryoForge targets local repair after the generation of an initial cryo-EM atomic model. The central problem is no longer to reconstruct the entire structure from the density map, but to identify local regions that remain problematic, determine whether they contain enough evidence for correction and choose an appropriate intervention. CryoForge therefore treats candidate residue regions as the units of repair and evaluates every attempted change against the local structural evidence (Fig. 1).

**Figure 1.**
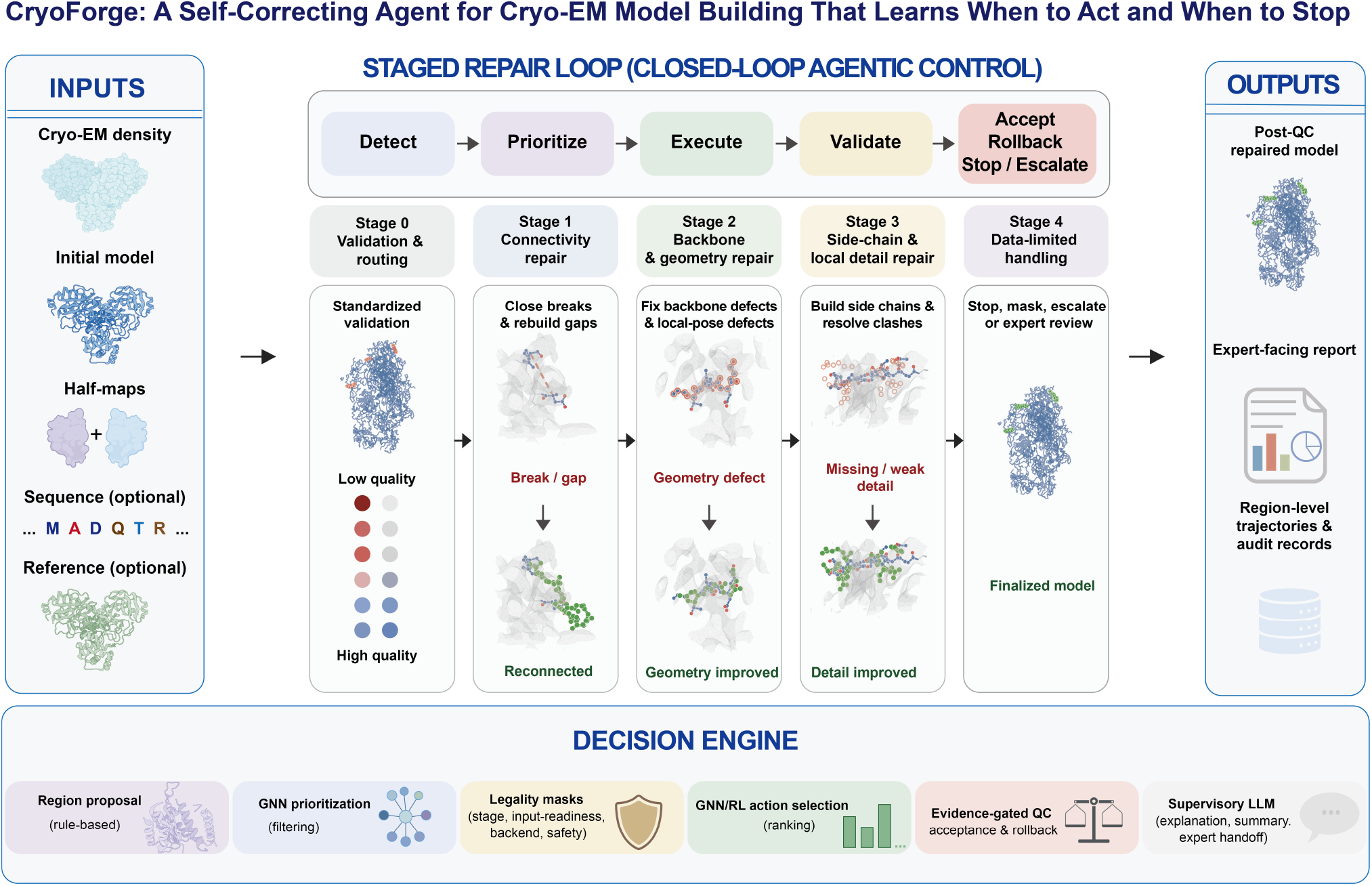
CryoForge overview: evidence-gated, learning-enhanced, region-level repair of cryo-EM initial models. CryoForge integrates cryo-EM density, initial coordinates, half maps, sequence information and an optional structural prior to identify candidate repair regions. Repair proceeds from connectivity and gaps to backbone conformation and side-chain detail. Learned models prioritize regions and repair choices, whereas post-repair QC determines whether an edit is accepted, reversed or stopped. A supervisory LLM summarizes the evidence and supports expert review without directly generating or editing atomic coordinates. Deposited models paired with benchmark entries are used only for offline label construction, benchmarking and retrospective evaluation.

CryoForge receives a cryo-EM density map, an initial atomic model, available half maps, sequence information and, when appropriate, an optional structural prior. It evaluates density fit, half-map consistency, local geometry, clashes, rotamers, Ramachandran conformation, peptide state, chain connectivity and neighboring structure at residue level. A sensitive rule-based screen then identifies local regions that may benefit from repair, providing candidates for subsequent learned prioritization.

The GNN estimates the repairability of each candidate from its local and neighboring evidence, and the GNN/RL policy ranks the available repair choices. Learned scores guide prioritization but do not determine acceptance: each attempted edit must improve its target defect without unacceptable loss of map support, stereochemical quality or neighboring structural stability. The supervisory LLM summarizes the evidence and supports interpretation of complex cases and expert handoff, but does not generate or edit atomic coordinates.

Local repair is organized from coarse to fine. Stage0 evaluates the input model; Stage1 addresses chain breaks, local gaps and continuity defects; Stage2 corrects backbone conformations, local poses and associated geometry defects; and Stage3 handles side-chain completion, rotamers, atomic clashes and local structural detail. Regions with insufficient evidence or non-unique interpretations are stopped or referred for expert review rather than forced into coordinate generation.

For each candidate region, CryoForge attempts the most appropriate stage-specific correction and then reassesses the resulting structure. Outcomes include validated improvement, low-confidence partial improvement, QC rejection with reversal, data-limited stopping, no effective gain and expert review. Only edits supported by the available evidence and stage-specific QC are incorporated into the repaired model.

The final output contains the repaired coordinates together with concise evidence summaries for expert inspection. An optional structural prior can assist local correspondence, whereas the deposited model paired with each benchmark entry is reserved for training and retrospective evaluation. In this way, CryoForge preserves the global structural framework while making local correction conservative and evidence dependent.

### Initial models retain a stage-structured local repair burden

To characterize the post-builder repair space, we assembled a resolution-stratified paired benchmark of 849 cryo-EM entries, with 283 entries in each of the 2.5–3.0, 3.0–3.5 and 3.5–4.0 Å ranges. Entries were split into 600 development, 120 validation and 129 held-out test entries. Each entry contributed a sequence-aware and a sequence-free or density-dominant initial model. Candidate-region behavior was characterized in the development cohort, with the validation and test partitions reserved for model selection and downstream evaluation, respectively. Benchmark construction and sample-mode definitions are described in Supplementary Methods 1.

Before quantifying repair burden, we asked whether the rule-based proposal captured the local defects identified by comparison with deposited models. It covered 93.5% of deposited-model-defined defect regions, with a residue-level recall of 93.0% (Fig. 2A). Matched candidate and defect regions had a mean intersection-over-union of 40.9% and a median boundary deviation of 2.2 residues (Fig. 2A,B). Thus, the proposal step captured most local defects while drawing moderately broader boundaries, as expected for a sensitive screen rather than a final segmentation model. Deposited models were used only for this benchmark comparison. Evaluation procedures are described in Supplementary Methods 2.

**Figure 2.**
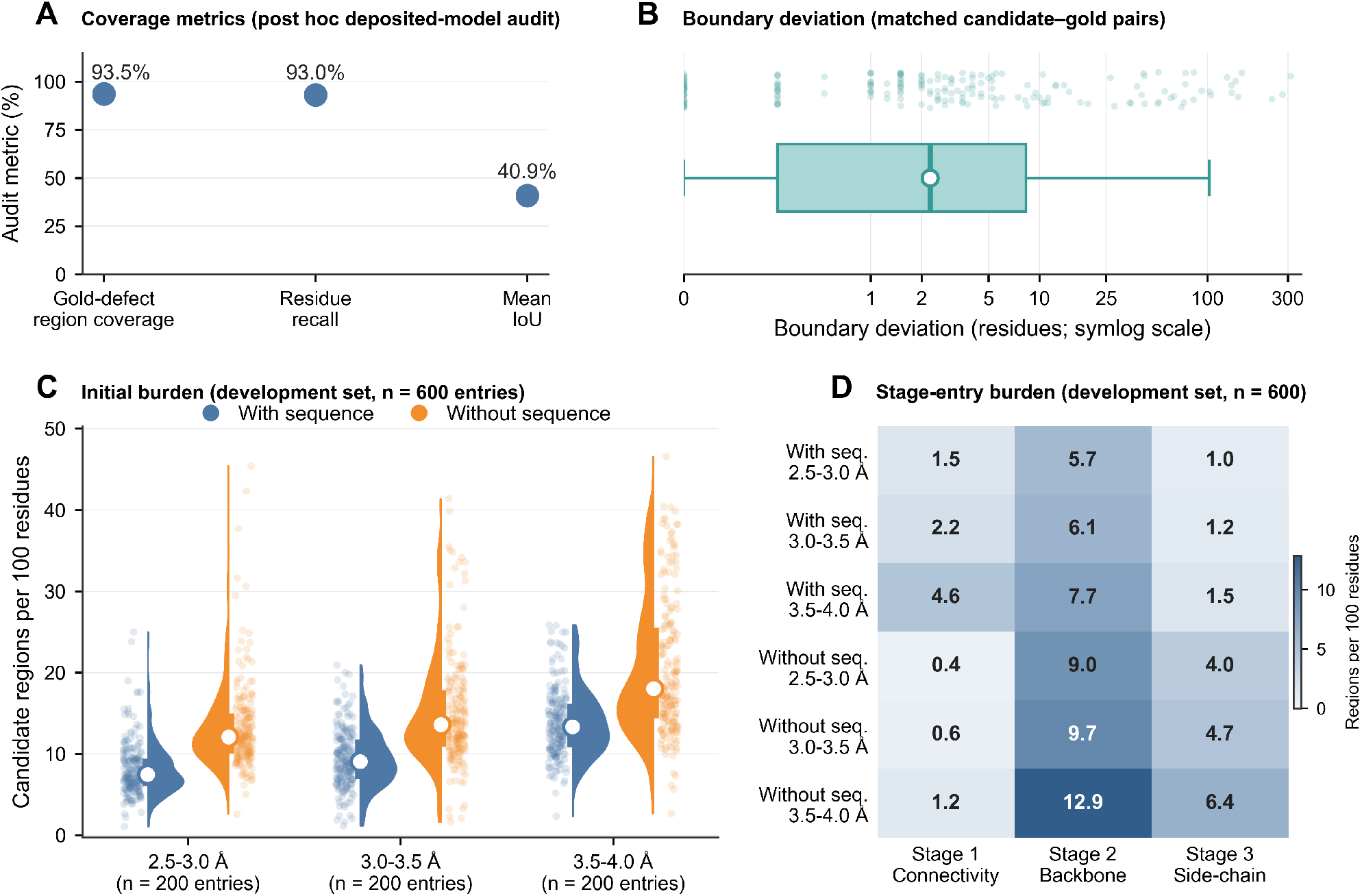
Cryo-EM initial models retain a systematic, stage-structured local repair burden. Candidate-region behavior was evaluated in the 600-entry development cohort. (A) Comparison with deposited models in eligible sequence-aware samples, reporting defect-region coverage, residue-level recall and mean intersection-over-union. Defect-region coverage is the fraction of deposited-model-defined defect regions overlapped by at least one same-chain candidate region. (B) Boundary deviation between matched candidate and defect regions. The white circle marks the median, the box-and-whisker plot summarizes the distribution and points denote matched intervals; the horizontal axis uses a symmetric-log scale. (C) Overall initial candidate-region burden per 100 modeled residues across all 1,200 paired sample modes, stratified by reported resolution and initialization mode. White circles denote medians, thick bars denote interquartile ranges and points denote individual sample modes. (D) Macro mean Stage1 connectivity, Stage2 backbone and Stage3 side-chain stage-entry burden in the same cohort. Candidate regions were re-detected on the model entering each stage and should not be summed across stages. Deposited models were used only for the comparisons in panels A and B.

We next quantified overall initial candidate-region burden in the development cohort. Across all three resolution strata, sequence-free or density-dominant initial models had a higher median burden than their paired sequence-aware models (Fig. 2C). As map resolution decreased, both initialization modes shifted toward higher burden and developed longer high-value tails. Sequence information therefore reduced but did not eliminate the candidate repair space, whereas lower resolution increased both typical burden and between-sample heterogeneity.

Residual burden was also distributed unevenly across repair levels (Fig. 2D). Stage2 backbone burden was highest in both initialization modes and every resolution stratum, increasing from 5.7 to 7.7 regions per 100 modeled residues in sequence-aware models and from 9.0 to 12.9 in sequence-free or density-dominant models as resolution decreased. Under the current stage-entry definition, sequence-free models had a lower Stage1 candidate burden but higher Stage2 and Stage3 burdens than sequence-aware models. This pattern does not indicate better overall connectivity; rather, it shows that their residual candidate space was concentrated more strongly in downstream backbone and side-chain repair. Because candidate regions were re-detected on the current model entering each stage, stage-specific burdens should not be summed.

The stage-specific compositions further supported this task stratification (Table 1). Stage1 was dominated by backbone-tracing defects, Stage2 comprised gap-insert pose and short-torsion defects, and Stage3 concentrated on side-chain rotamers, clash or pose defects and other local side-chain interpretation problems. Stage1–Stage3 therefore represent distinct repair tasks rather than repeated classification of one defect class. Together, these analyses show that post-builder repair burden is structured by map resolution, sequence availability and defect type, motivating separate connectivity, backbone and side-chain repair stages.

**Table 1.**
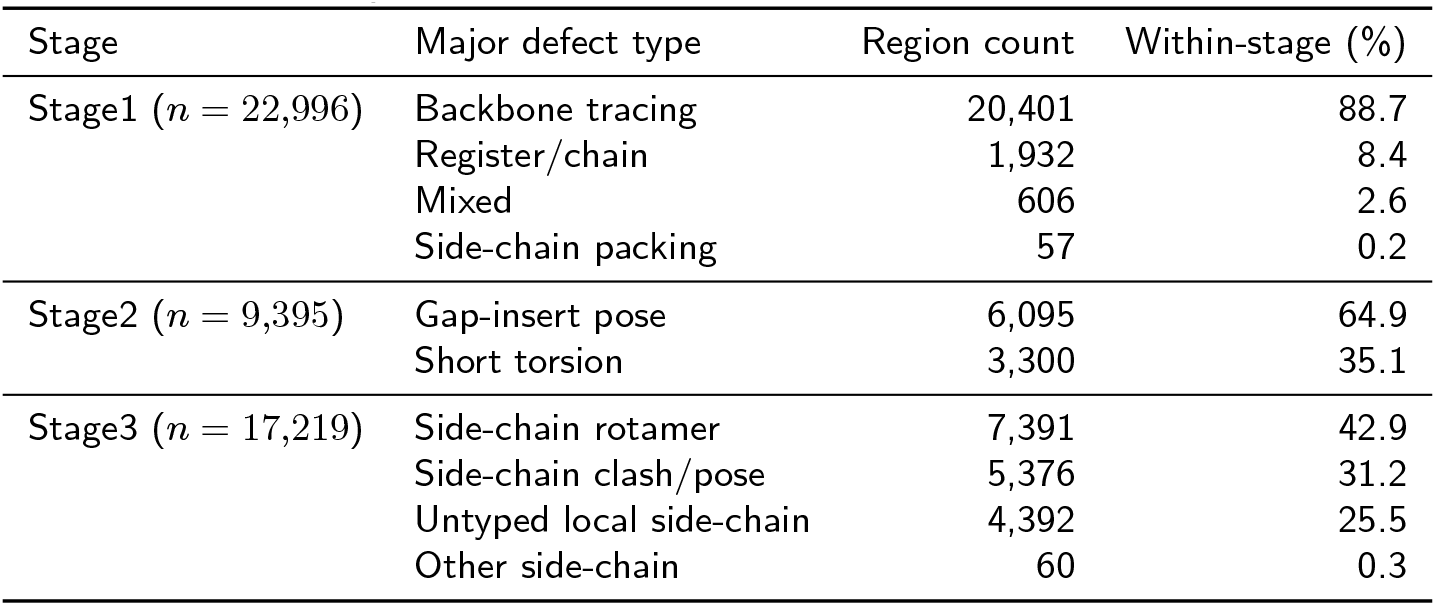
Stage-specific candidate-region composition in the development cohort. Candidate regions were re-detected on the current model entering each stage; counts therefore represent distinct stage-entry candidate sets and should not be summed across stages.

### Staged closed-loop repair converts candidate regions into validated local improvements

After identifying candidate regions in the initial models, we evaluated whether CryoForge could convert them into verifiable local structural improvements in the held-out full-QC sequence-aware test cohort. Regions with sufficient initial evidence entered staged repair: gaps and connectivity defects were addressed first, followed by local backbone or pose errors and then side-chain completion, rotamer correction and clash-aware refinement. The choice of repair depended on the local defect and its supporting evidence, and every attempted change was reassessed before it could enter the final model.

The overall workflow summarized in Fig. 1 was applied here as a coarse-to-fine repair trajectory. CryoForge first addressed connectivity and gap defects that could affect downstream interpretation in Stage1, then corrected local backbone geometry and pose in Stage2, and finally performed atom-level completion, side-chain placement and local stereochemical refinement in Stage3. After each attempted edit, a stage-specific QC gate determined whether the candidate state was accepted, rolled back, stopped or marked as data-limited. Thus, Stage1, Stage2 and Stage3 formed a staged trajectory rather than three independent parallel repair tasks.

To illustrate this closed-loop process in a concrete structural context, we selected a standard validated-improved multi-stage trajectory and visualized the initial model, stage-specific local repair windows and final post-QC model (Fig. 3A). This sample contained multiple residual defects that required different repair operations rather than a single isolated cleanup step. Stage1 reconnected a local gap, Stage2 corrected the local backbone conformation and Stage3 completed the remaining side-chain detail. The final model retained the surrounding structural context while incorporating only the local edits that passed the corresponding stage-specific QC gates. This trajectory illustrates the intended coarse-to-fine progression from connectivity repair through backbone cleanup to atom-level completion.

**Figure 3.**
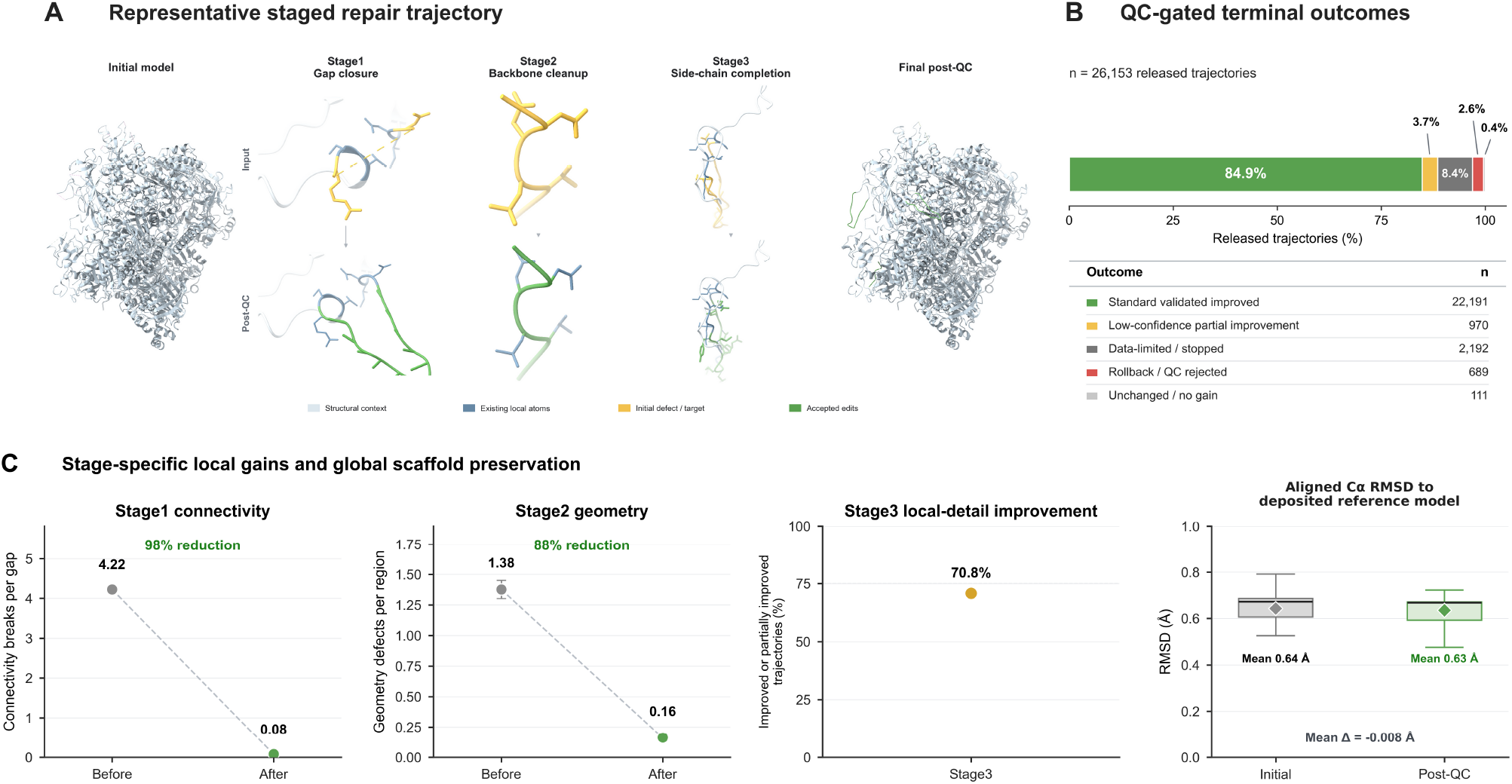
Staged QC-gated repair produces local structural gains while preserving the global scaffold. (A) Representative standard validated-improved trajectory shown from the initial model through Stage1 gap closure, Stage2 backbone cleanup and Stage3 side-chain completion to the final post-QC model. Pale blue denotes structural context, dark blue denotes existing local atoms, yellow denotes the initial defect or target and green denotes accepted edits. The example was selected for visual interpretability and was not used to estimate aggregate performance. (B) Terminal outcomes across 26,153 released repair trajectories in the held-out full-QC sequence-aware test cohort. Standard validated improvements accounted for 22,191 trajectories (84.9%), and caution-labeled low-confidence partial improvements accounted for 970 (3.7%). Together, these classes form the aggregate validated-improved group used in the learned-decision analyses. The remaining outcomes were data-limited or stopped, rollback-protected or QC-rejected, and unchanged or no gain; no worsened trajectory was retained. (C) Stage-specific local gains and global-scaffold preservation in the same cohort. Mean Stage1 connectivity breaks per gap decreased from 4.22 to 0.08, and mean Stage2 geometry defects per region decreased from 1.38 to 0.16. Among evaluated Stage3 local-detail edits, 70.8% achieved a local-detail gain. Boxplots summarize aligned Cα RMSD to the deposited model before and after QC; diamonds denote means of 0.64 and 0.63 Å, respectively, with a mean paired change of *−* 0.008 Å. Deposited models were used only for retrospective comparison.

Across all 26,153 released repair trajectories in the held-out full-QC sequence-aware test cohort, 22,191 (84.9%) were standard validated improvements and 970 (3.7%) were caution-labeled low-confidence partial improvements that passed all non-regression safeguards and were promoted (Fig. 3B). Together, these two classes comprised 23,161 trajectories (88.6%) and define the aggregate validated-improved class used in the learned-decision analyses. Of the remaining trajectories, 2,192 (8.4%) were stopped as data-limited, 689 (2.6%) were rejected by QC with rollback protection and 111 (0.4%) terminated unchanged or without effective gain. No worsened trajectory was retained in the final promoted outputs (*n* = 0).

Aggregate stage-specific summaries confirmed that the repair stages acted on their intended local targets (Fig. 3C). Mean Stage1 connectivity breaks per gap decreased from 4.22 before repair to 0.08 after QC, corresponding to a 98% reduction. Mean Stage2 geometry defects per region decreased from 1.38 to 0.16, an 88% reduction. Among evaluated Stage3 local-detail edits, 70.8% achieved a local-detail gain, comprising standard validated improvements and caution-labeled low-confidence partial improvements.

Together, these summaries show stage-specific gains in connectivity, local geometry and side-chain or local-detail interpretation.

Mean aligned Cα RMSD to the deposited reference model remained nearly unchanged, decreasing from 0.64 Å in the initial models to 0.63 Å after QC, with a mean paired change of *−*0.008 Å (Fig. 3C). Thus, substantial stage-specific local gains were achieved without materially altering global model-to-reference agreement, supporting the intended scope of CryoForge as a local repair system rather than a whole-structure rebuilding method.

Together, these results show that CryoForge converts candidate regions into validated local improvements through staged, QC-gated repair rather than unconstrained global rebuilding. Evidence-supported edits were retained, unsafe attempts were rejected or reversed and data-limited regions were left unchanged, with no worsened result entering the final model.

### Non-improving events reveal stage-specific evidence boundaries for automatic local repair

The preceding results showed that CryoForge converted most released repair trajectories into QC-supported local improvements. Automatic repair should not, however, force a structural change into every candidate region. When local evidence is insufficient or conflicting, the safer outcome is to stop, reject or reverse an attempted edit. We therefore examined the regions that did not yield a validated improvement at each stage. Because regions were reassessed as the model changed, the stage-specific event sets were analyzed separately and were not pooled.

Boundary composition differed sharply by stage (Table 2). Among 834 Stage1 events, weak or discontinuous density was most common (61.9%), followed by long gaps or large missing segments (27.7%). Stage2 was dominated by geometry–map tradeoffs, which accounted for 89.4% of 1,819 non-improving events. Stage3 boundaries were more varied: weak or discontinuous density accounted for 39.5% of 1,249 events, absence of a safe repair for 34.6%, insufficient structural support for 13.9% and sequence/register ambiguity for 8.0%. Thus, connectivity repair was limited mainly by density and gap feasibility, backbone repair by geometry–map conflict and side-chain repair by a broader combination of weak evidence and structural ambiguity.

**Table 2.** Stage-specific composition of evidence-boundary events. Percentages were calculated within each stage-specific event set. Candidate regions were re-evaluated at stage entry; therefore, event counts are not additive across stages.

| Stage | Event set | *n* | Long gap /<br>large missing segment | Weak or<br>discontinuous density | Geometry-map<br>tradeoff | No safe<br>repair | Insufficient<br>structural support | Sequence/register<br>ambiguity |
| --- | --- | --- | --- | --- | --- | --- | --- | --- |
| Stage1 | Triage-boundary events | 834 | 231 (27.7%) | 516 (61.9%) | — | — | 87 (10.4%) | — |
| Stage2 | Non-improving events | 1,819 | 40 (2.2%) | 144 (7.9%) | 1,627 (89.4%) | 8 (0.4%) | — | — |
| Stage3 | Non-improving events | 1,249 | 51 (4.1%) | 493 (39.5%) | — | 432 (34.6%) | 173 (13.9%) | 100 (8.0%) |

To illustrate these stage-specific boundary classes, we examined three representative trajectories (Fig. 4A–C). These examples were not treated as random software failures. Instead, they corresponded to three common limits of automatic repair: a data-limited stop caused by a weak-density long gap, a no-effective-gain outcome caused by geometry–map tradeoff and a QC rejection or rollback-protected outcome caused by unsupported side-chain completion.

**Figure 4.**
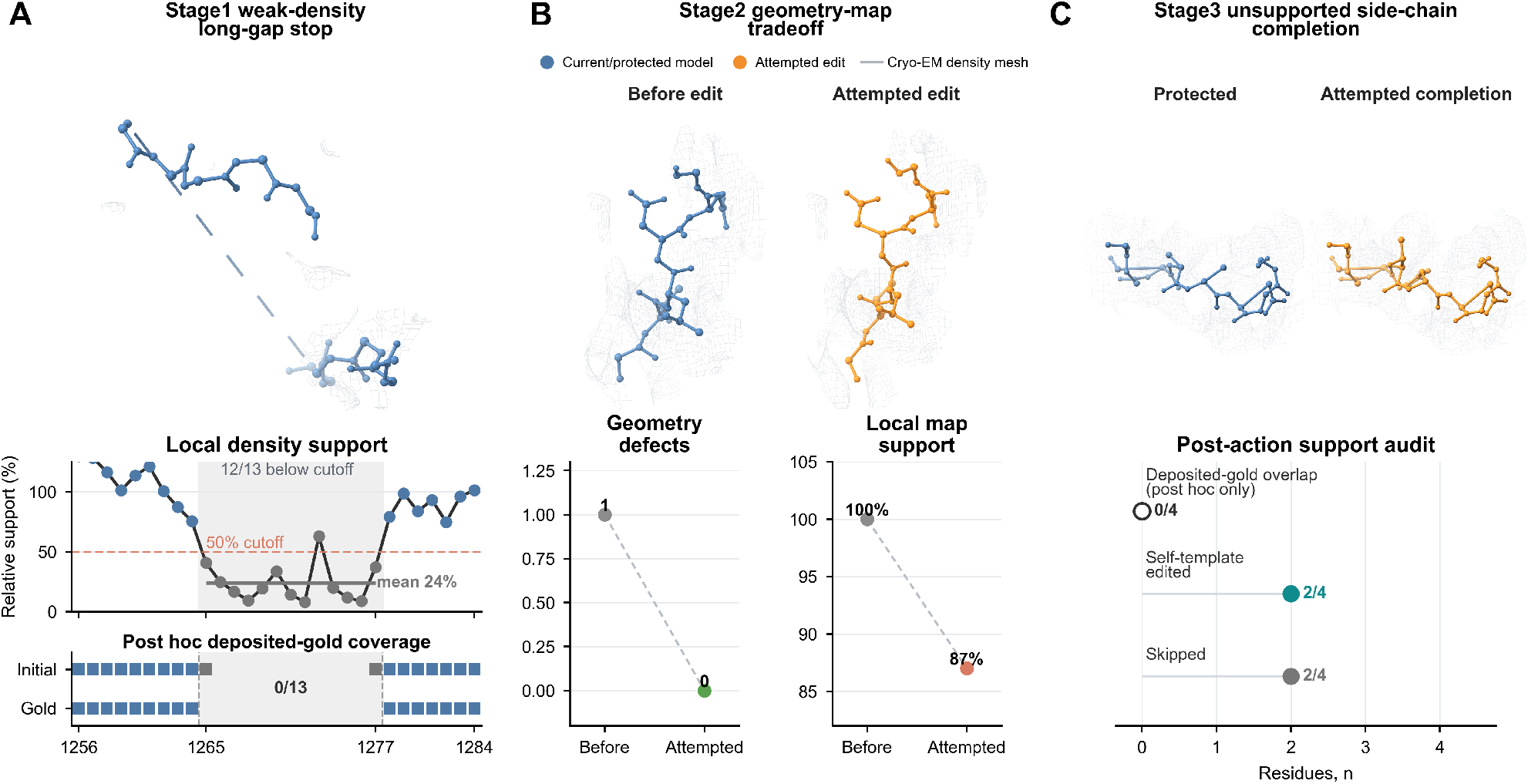
Representative evidence boundaries for automatic local repair. (A) Stage1 weak-density long-gap stop spanning residues 1265–1277. Twelve of thirteen gap residues fell below the 50% anchor-normalized density cutoff and mean gap support was 24%, leading to a data-limited stop based on runtime density evidence. Post hoc audit additionally showed that the deposited reference model contained no corresponding residues in the interval. (B) Stage2 geometry–map tradeoff. The attempted edit reduced geometry defects from 1 to 0 but decreased local map support from 100% to 87%, producing an unchanged or no-effective-gain outcome rather than promotion into the accepted model. (C) Stage3 unsupported side-chain completion in Chain Ba residues 886–889. The attempted edit completed 2 of 4 residues and skipped 2 of 4; none of the four positions overlapped the deposited model. The attempted completion was rejected by post-repair QC and the protected coordinates were retained. Blue denotes the current or protected model, orange denotes the attempted edit and grey mesh denotes cryo-EM density. Deposited models are shown only for retrospective comparison.

The first boundary case was a weak-density long-gap region (Fig. 4A). The initial model contained a gap spanning residues 1265–1277. Anchor residues on both sides retained local density support, but support decreased sharply inside the gap: 12 of 13 gap residues fell below the 50% cutoff, and mean support was only 24% of that at the anchors. CryoForge therefore stopped rather than forcing completion of a structure unsupported by the runtime density evidence. Post hoc audit additionally showed that the deposited reference model contained no corresponding residues in this interval. This case illustrates how weak local evidence can define a genuine limit of automatic repair.

The second boundary case represented a geometry–map tradeoff (Fig. 4B). The attempted edit reduced local geometry defects from 1 to 0, indicating a genuine improvement in stereochemical terms. At the same time, however, local map support decreased from 100% to 87%. Because validated repair required local geometry improvement without map-support regression beyond the QC tolerance, this trajectory was not accepted as a validated improvement and was instead classified as unchanged or no effective gain. Thus, some regions were not completely uneditable; rather, the available edit failed to produce a net QC-supported gain.

The third boundary case involved unsupported side-chain completion (Fig. 4C). The attempted correction completed only 2 of 4 residues in Chain Ba residues 886–889, while the remaining two were skipped, and none of the four positions overlapped the deposited model. Post-repair QC found insufficient support for the edited structure, so the attempted completion was rejected and the protected coordinates were retained. This example shows that a technically possible edit is not necessarily a structurally justified one.

Together, these analyses show that stage-specific non-improving events mainly reflect limits in structural evidence rather than arbitrary software failure. Weak density, geometry–map tradeoffs, sequence or register ambiguity and the absence of a safe correction led CryoForge to stop, reject or reverse an edit. These conservative outcomes help prevent over-modeling and keep unsupported changes out of the final model.

### Learned decision layers improve candidate filtering and repair-action selection

In CryoForge, the rule-based candidate-region proposal module was designed to provide a high-sensitivity candidate pool. Its purpose was to reduce missed repairable regions rather than to define a high-precision final repair target set. This design preserved most potentially actionable regions, but it also introduced many candidates that did not subsequently enter the aggregate validated-improved class. We therefore evaluated whether learned decision layers could improve downstream repair decisions on the same held-out full-QC sequence-aware test candidate-region pool, legal action space, QC gates and rollback mechanism. We compared a Rule-only strategy, a GNN-only learned gate and the full GNN/RL strategy.

We first asked whether learned gates could reduce non-improving candidate burden while preserving high recall. In these analyses, recall measures the fraction of regions in the aggregate validated-improved class that were retained, whereas precision measures the fraction of retained candidates that entered this class after QC. Promoted low-confidence partial-improvement subtypes were included in the aggregate positive class because they passed all non-regression safeguards, although their caution labels remained available. In Stage2, Rule-only priority had low precision when recall was kept high, indicating that many additional candidates had to be retained to avoid missing repairable regions. Both GNN-only and GNN/RL learned gates increased precision across the same recall range (Fig. 5A). Thus, the learned models did not replace rule-based candidate proposal. Instead, they acted on top of the high-sensitivity rule-generated pool to prioritize candidate regions more likely to be actionable repair targets.

**Figure 5.**
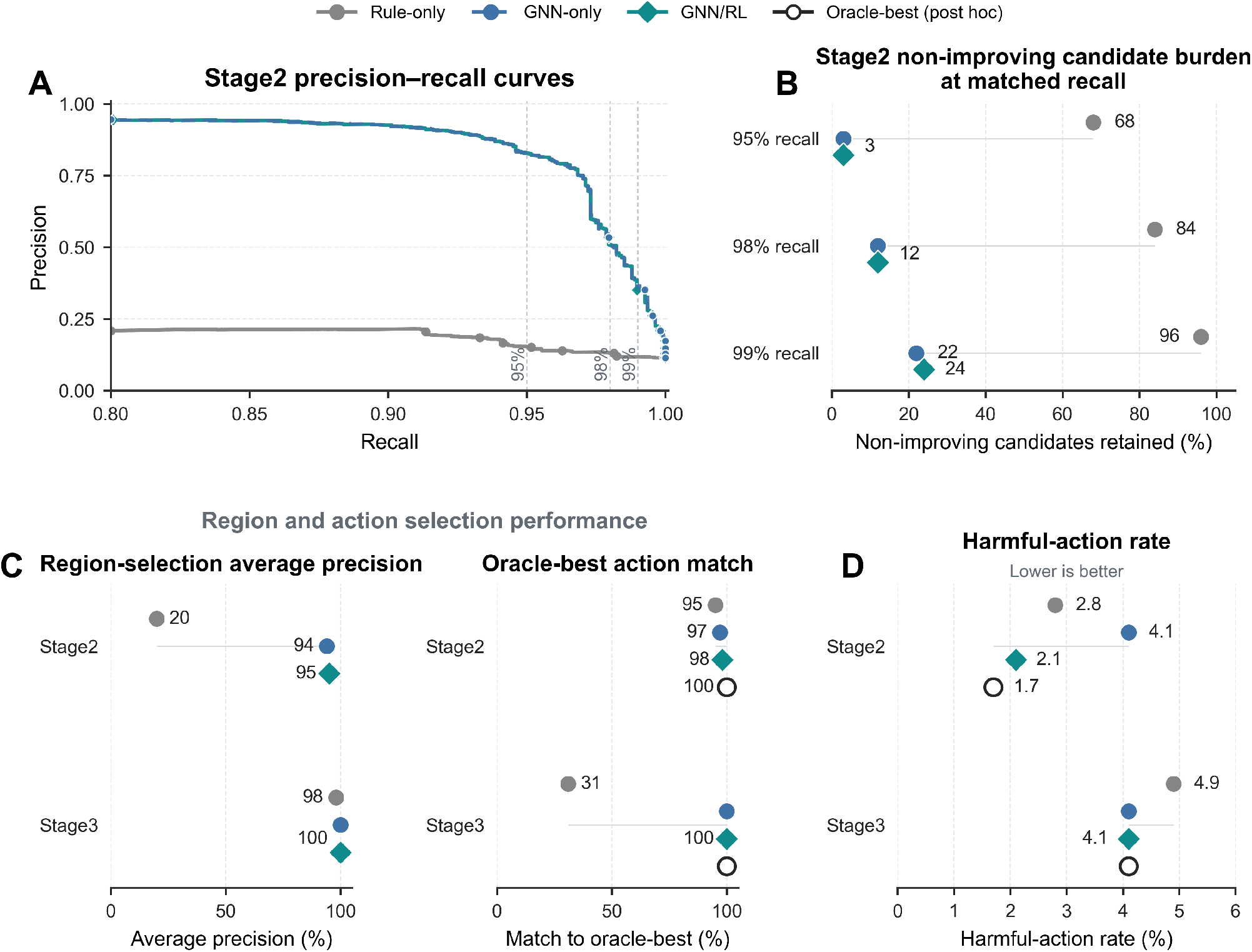
Learned decision layers improve candidate-region filtering and repair-action selection. (A) Stage2 precision–recall curves comparing Rule-only priority, GNN-only filtering and the full GNN/RL strategy in the held-out full-QC sequence-aware test cohort. The GNN-only and GNN/RL curves largely overlap. (B) Percentage of non-improving candidates retained at matched validated-improved recall of 95%, 98% and 99%. Promoted low-confidence partial improvements are included in the positive class. (C) Region- and action-selection performance. The left panel reports region-selection average precision in Stage2 and Stage3; the right panel reports agreement with the best-performing available action identified retrospectively for regions with multiple choices. After learned region filtering, GNN-only ranks legal actions by predicted immediate gain, whereas GNN/RL ranks the same legal actions by predicted Q. (D) Harmful-action rate for Rule-only, GNN-only, GNN/RL and the retrospective best-action reference in Stage2 and Stage3; lower values indicate safer action selection. All strategies were evaluated on the same held-out candidates under the same QC and rollback criteria. Definitions are provided in Supplementary Methods 4 and Supplementary Methods 5.

The reduction in non-improving candidate burden was clearer at fixed recall operating points. When validated-improved recall was constrained to 95%, 98% and 99%, the Rule-only strategy retained 68%, 84% and 96% of non-improving candidates, respectively. GNN-only retained 3%, 12% and 22%, while GNN/RL retained 3%, 12% and 24% at the same recall levels (Fig. 5B). The near-identical learned-strategy profiles indicate that this filtering benefit arose primarily from the GNN-based learned gate rather than from the downstream RL action policy.

Average precision provided a threshold-independent summary of the same region-filtering behavior. In Stage2, region-selection average precision increased from 20% with Rule-only priority to 94% with GNN-only and 95% with GNN/RL. Stage3 began near ceiling at 98% with Rule-only and reached 100% with both learned strategies (Fig. 5C, left). Thus, the learned gate produced its largest region-selection gain in the more heterogeneous Stage2 candidate pool, whereas the Stage3 pool was already comparatively clean.

We next evaluated action selection for candidate regions with multiple possible repairs, using the retrospectively identified best-performing action as an upper-bound reference. GNN-only ranked legal actions by predicted immediate gain, whereas GNN/RL ranked the same available choices by predicted Q, incorporating expected gain, risk and downstream utility. All methods were compared on the same multi-action decision sets (Stage2, *n* = 9,106; Stage3, *n* = 16,770). In Stage2, agreement with the best-performing action increased from 95% with Rule-only selection to 97% with GNN-only and 98% with GNN/RL. In Stage3, agreement was 31% for Rule-only and 100% for GNN-only and GNN/RL (Fig. 5C, right). Thus, learned immediate-gain ranking accounted for most of the action-selection improvement, while Q-based ranking provided an additional risk-aware distinction in Stage2.

Finally, harmful-action analysis showed that GNN/RL provided additional action-level safety. In Stage2, its harmful-action rate was 2.1%, lower than Rule-only (2.8%) and GNN-only (4.1%) and close to the retrospective best-action reference (1.7%). In Stage3, GNN/RL and GNN-only each had a harmful-action rate of 4.1%, below Rule-only at 4.9% (Fig. 5D). The additional value of the RL/Q-policy was therefore most evident in Stage2, where region filtering alone did not minimize harmful choices. Stage3 was already near ceiling for the learned strategies, leaving less room for improvement.

Together, these results support the layered CryoForge design. Rule-based proposal provides sensitive coverage, the GNN improves identification of likely repairable regions and the GNN/RL policy contributes risk-aware selection among repair choices. The learned components reduced non-improving candidate burden while preserving recall and reduced harmful choices without weakening structural QC.

### CryoForge distinguishes underbuilt and overbuilt-like residual-error phenotypes across initialization regimes

Finally, we asked whether CryoForge could operate downstream of a contemporary AI-based cryo-EM model builder and whether its residual errors differed from those of the main-benchmark initialization workflow, hereafter termed the benchmark workflow. We selected CryoAtom as the external initializer because it represents a technically distinct map-to-model strategy [14]. Using the same 129 held-out test entries for both initializations, we compared their initial structures, characterized the candidate regions identified by CryoForge and applied the same conservative repair and QC procedure to the CryoAtom-derived models. Cα-level comparison methods are described in Supplementary Methods 1. Deposited models were used only for retrospective characterization.

The two initialization regimes produced clearly different residual-error profiles (Fig. 6A). CryoAtom-derived models had substantially better backbone continuity than benchmark-workflow initial models, with Cα breaks decreasing from 21.7 to 0.8 breaks per 1000 Cα atoms. They also covered a slightly larger fraction of deposited-model Cα atoms than benchmark-workflow initial models (81.6% versus 77.4%). However, the fraction of modeled Cα atoms that could be matched to the deposited model was lower for CryoAtom-derived models than for benchmark-workflow initial models (60.7% versus 73.2%), and the unmatched modeled Cα fraction was correspondingly higher (39.3% versus 26.8%). Because these matched and unmatched fractions are computed over modeled Cα atoms, this pattern indicates that CryoAtom did not simply reduce the same residual errors observed in the benchmark workflow. Instead, the dominant residual phenotype shifted from fragmented or underbuilt models toward more continuous but scope-expanded or overbuilt-like regions in the retrospective comparison.

**Figure 6.**
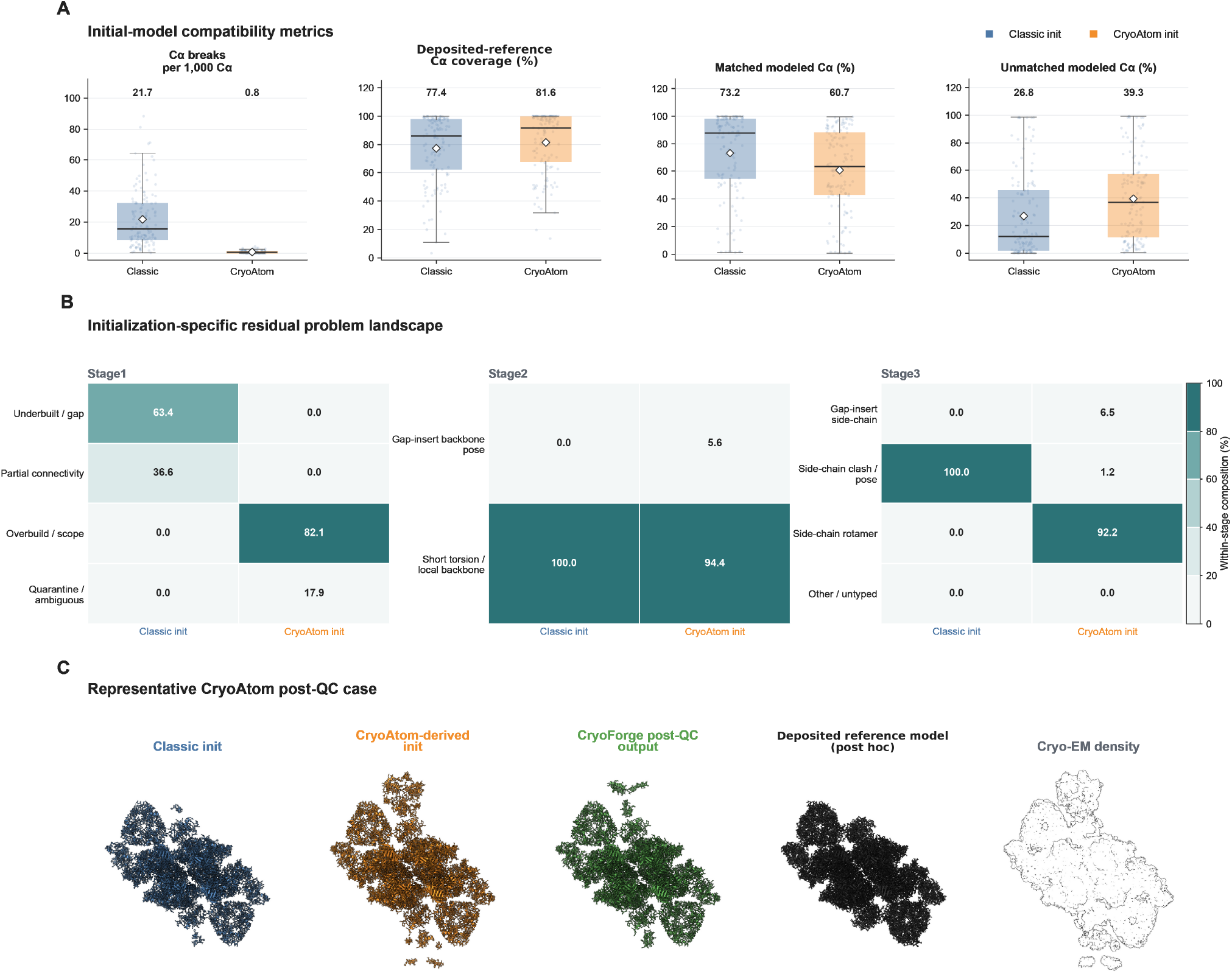
CryoForge distinguishes underbuilt and overbuilt-like residual-error phenotypes across initialization regimes. (A) Initial-model comparison between benchmark-workflow and CryoAtom-derived models for the same 129 held-out test entries. CryoAtom-derived models showed fewer C*α* breaks and slightly higher deposited-model C*α* coverage, but lower matched and higher unmatched modeled C*α* fractions. Coverage is expressed relative to deposited-model C*α* atoms, whereas matched and unmatched fractions are expressed relative to modeled C*α* atoms. (B) Candidate-region composition for the two initialization methods. Each heat-map column is normalized within its stage and initialization and describes the residual-error profile rather than repair performance. (C) Representative subtractive CryoAtom case, showing the benchmark-workflow initial model from the same entry, the CryoAtom-derived input, the post-QC output, the deposited model and the cryo-EM density. Unsupported CryoAtom-derived coordinates were removed or withheld rather than extended. Phenotype definitions are provided in Supplementary Methods 2.

The matched 129-entry comparison also revealed initialization-specific stage-entry candidate compositions (Fig. 6B). Percentages were calculated separately within each initialization and stage, so the panel describes the composition of the candidate pools rather than their absolute burden or repair-method performance. In Stage1, classic candidates comprised 63.4% underbuilt or gap-like regions and 36.6% partial-connectivity regions, whereas CryoAtom candidates comprised 82.1% overbuilt or scope-expanded regions and 17.9% quarantine or ambiguous regions. Stage2 classic candidates were entirely short-torsion or local-backbone regions, while CryoAtom candidates were predominantly short-torsion or local-backbone regions (94.4%) with a smaller gap-insert backbone-pose component (5.6%). Stage3 classic candidates were entirely side-chain clash or pose regions; CryoAtom candidates were dominated by side-chain rotamers (92.2%), with smaller gap-insert side-chain (6.5%) and side-chain clash or pose (1.2%) components. Thus, the same held-out entries generated different residual candidate phenotypes under the two initialization routes.

CryoForge identified 8,094 Stage1, 7,288 Stage2 and 400 Stage3 candidate regions in the CryoAtom-derived models. Most lacked sufficient evidence or a sufficiently safe correction for automatic repair. Only six Stage2 regions and nine Stage3 regions proceeded to an attempted edit, illustrating the deliberately conservative treatment of scope-expanded and weakly supported models.

Terminal outcomes were summarized using stage-appropriate analysis units (Table 3). At Stage1, one sample-level terminal outcome was assigned to each held-out entry: 35 of 129 entries (27.1%) were standard validated improved, 75

**Table 3.** Stage-specific outcomes in the CryoAtom held-out compatibility analysis. Percentages were calculated within each row. Stage1 reports one sample-level outcome for each of the 129 held-out entries, whereas Stage2 and Stage3 report repaired candidate regions.

| Stage | Analysis unit | Total | Standard validated improved | Low-confidence partial improvement | Manual review | Conservatively protected / no action | Rollback-protected / QC-rejected |
| --- | --- | --- | --- | --- | --- | --- | --- |
| Stage1 | Held-out entry, sample-level terminal outcome | 129 | 35 (27.1%) | 0 | 75 (58.1%) | 19 (14.7%) | 0 |
| Stage2 | Released region trajectory | 6 | 5 (83.3%) | 0 | 0 | 0 | 1 (16.7%) |
| Stage3 | Released region trajectory | 9 | 0 | 9 (100.0%) | 0 | 0 | 0 |

No data-limited, unchanged or worsened outcomes occurred in these three stage-specific groups.

(58.1%) were directed to manual review and 19 (14.7%) were conservatively protected without an automatic action. Among the six released Stage2 region trajectories, five (83.3%) were standard validated improved and one (16.7%) was rollback-protected or QC-rejected. All nine released Stage3 trajectories were retained as caution-labeled low-confidence partial improvements. No data-limited or stopped, unchanged or no-effective-gain, or final-worsened-retained outcome occurred in these stage-specific external-initialization summaries.

The representative case in Fig. 6C illustrates the mainly subtractive role of post-builder handling for a scope-expanded CryoAtom model. CryoForge removed or withheld unsupported coordinates to produce a more conservative post-QC structure rather than extending the model further. Together, these results show that different initialization methods leave different kinds of residual errors and therefore benefit from different downstream responses: completion-oriented repair for fragmented models, but more frequent withholding or removal for continuous yet weakly supported regions.

### Challenge-derived case studies define the practical division of labor between CryoForge and expert-assisted modeling

To assess how CryoForge could contribute in expert-facing model-repair settings, we selected three Team 41 cases from the 2019 EMDataResource Model Challenge and its archived dataset [5, 30], spanning 1.8–3.1 Å, different protein sizes and distinct initial-model quality regimes. Each case contained fully automated and manual-assisted submissions from the same target and team, with approximately 1 h of reported human effort for the manual-assisted submission. The automated submission was used as the CryoForge input, whereas the HUMAN submission and deposited reference model were reserved for retrospective behavioral comparison and quantitative audit. None of the three challenge targets occurred in the frozen 849-entry benchmark or learned-policy training data. Because the challenge archive did not retain exact parent-submission identifiers or chronological expert-operation logs, AUTO-to-HUMAN coordinate differences were interpreted as post hoc expert-edit footprints rather than literal operation histories. The resulting models and quantitative comparisons are summarized in Fig. 7 and Table 4.

**Table 4.** Repair scope and selected quality metrics in the challenge-derived cases. CryoForge edit counts are QC-promoted action footprints. HUMAN backbone and side-chain counts are post hoc AUTO-to-HUMAN coordinate-difference phenotypes rather than logged expert operations. Metric pairs are CryoForge/HUMAN; deposited-model metrics are retrospective.

| Case | Repair footprint<br>CryoForge / HUMAN | Mean<br>Q-score | Clashscore | Ramachandran<br>outliers (%) | Rotamer<br>outliers (%) | $C\alpha$ RMSD<br>(Å) | GDT<sub>HA</sub><br>(%) | $C\alpha$-<br>IDDT |
| --- | --- | --- | --- | --- | --- | --- | --- | --- |
| T0101, 1.8 Å | CryoForge: removed A177; 0 BB; 0 detail | 0.865/0.870 | 2.89/4.67 | 1.18/0.58 | 3.27/2.60 | 0.284/0.309 | 98.55/98.26 | 0.994/0.992 |
|  | HUMAN: 1 BB; 2 SC; 0 cofactors |  |  |  |  |  |  |  |
| T0104, 2.9 Å | CryoForge: +10 residues; 3 BB; 26 detail | 0.688/0.698 | 33.34/9.99 | 4.98/0.81 | 7.81/17.53 | 0.463/0.455 | 94.08/94.92 | 0.966/0.967 |
|  | HUMAN: +10 residues; 44 BB; 187 SC; +2 NAD |  |  |  |  |  |  |  |
| T0103, 3.1 Å | CryoForge: +2 residues; 3 BB; 6 detail | 0.641/0.657 | 54.45/12.00 | 12.43/0.60 | 17.65/16.45 | 0.443/0.570 | 93.60/89.41 | 0.959/0.937 |
|  | HUMAN: 0 residue-set change; 26 BB; 57 SC |  |  |  |  |  |  |  |

BB, backbone footprint residues; SC, side-chain footprint residues. For T0104 and T0103, whole-model Q-score and MolProbity values use the standardized QC snapshot immediately before the final low-confidence terminal extension; the final A1 or A176 extension was accepted by local anchor, density or half-map proxy, geometry and continuity QC. Deposited-reference coordinate metrics use the displayed final models.

**Figure 7.**
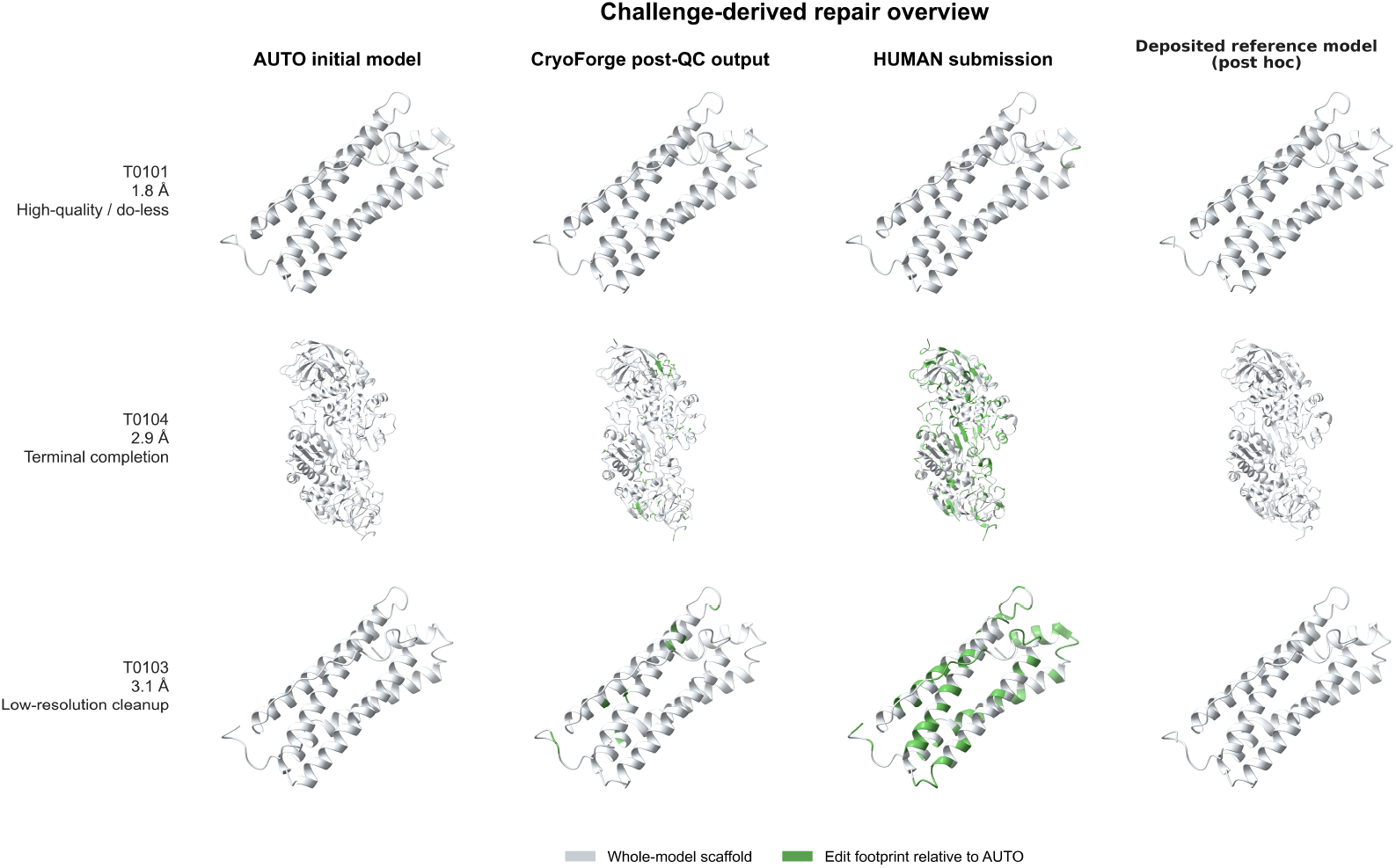
Challenge-derived case studies reveal complementary repair footprints of CryoForge and expert-assisted modeling. Rows show T0101 at 1.8 Å, T0104 at 2.9 Å and T0103 at 3.1 Å. Columns show the AUTO initial model, CryoForge post-QC output, HUMAN submission and deposited reference model used only for post hoc audit. Whole-model scaffolds are light grey. Green regions denote QC-promoted CryoForge actions or retrospective AUTO-to-HUMAN coordinate-edit footprints based on residue-set changes, C*α* displacement of at least 0.5 Å or side-chain RMSD of at least 1.0 Å. HUMAN footprints are coordinate-difference phenotypes, not chronological operation logs. Neither the HUMAN submission nor the deposited reference model was available to CryoForge at runtime.

In the 1.8 Å T0101 apoferritin case, the AUTO model was already highly complete and locally well resolved. CryoForge removed only A177, which lay outside the supplied target-sequence scope, and promoted no additional backbone or side-chain repairs. The HUMAN submission retained the same residue set and differed at only one backbone and two side-chain footprint residues. Both outputs were close to the deposited reference model. CryoForge retained a lower clashscore and C*α* RMSD, whereas HUMAN had fewer Ramachandran, rotamer and C*β* outliers. Thus, when the starting model was already close to acceptable, CryoForge performed a defined scope correction without introducing broader changes.

In the 2.9 Å T0104 alcohol dehydrogenase case, CryoForge and HUMAN restored exactly the same ten terminal residues: A1, A371–374, B1 and B371–374. CryoForge additionally promoted local backbone repairs involving three residues and local-detail repairs involving 26 residues. In contrast, post hoc AUTO-to-HUMAN comparison identified 44 backbone and 187 side-chain footprint residues, together with two added NAD cofactors. The two workflows therefore agreed on the clearly bounded completeness task but differed substantially in their wider coordinate-edit scope.

This difference was also reflected in model-quality metrics. The T0104 HUMAN model had a higher mean Q-score, lower clashscore, fewer Ramachandran outliers and slightly higher GDT_HA and C*α*-lDDT. CryoForge nevertheless retained fewer rotamer outliers, while C*α* RMSD and C*α*-lDDT were nearly identical between the two outputs. Broad expert-assisted remodeling therefore improved overall stereochemistry without uniformly improving every local or coordinate-agreement metric.

In the 3.1 Å T0103 apoferritin case, CryoForge added A5 and A176 as low-confidence terminal completions and promoted local backbone repairs involving three residues and local-detail repairs involving six residues. The HUMAN submission did not change the residue set, but its coordinate-difference footprint covered 26 backbone and 57 side-chain residues. HUMAN remodeling markedly reduced clashscore and Ramachandran outliers and increased mean Q-score. However, CryoForge remained closer to the deposited reference model by C*α* RMSD, GDT_HA and C*α*-lDDT. This result shows that broad stereochemical cleanup and retrospective agreement with deposited coordinates are related but non-equivalent objectives.

CryoForge was more conservative in broad backbone editing because its promotion criterion required a verifiable net gain across local map or half-map support, geometry, connectivity and neighboring structural stability. Candidate regions required clear sequence boundaries, stable anchors, an interpretable density path or a legal and locally map-compatible structural prior. Even when an attempted edit improved a geometric metric, it was not promoted if map support regressed, neighboring structure was damaged, connectivity became less certain or the overall gain remained insufficient. The frozen action registry also focused on local gap closure, short-window backbone correction and side-chain detail completion rather than domain-scale repositioning, long-range register rebuilding, whole-model coupled refinement or cofactor interpretation.

Together, the three cases reveal complementary repair scopes rather than a single universally superior output. CryoForge reproduced selected expert-like completeness decisions, avoided unnecessary changes in an already highquality model and promoted a traceable subset of local edits. HUMAN submissions included broader backbone, side-chain and chemical-component remodeling, which substantially improved stereochemistry in the intermediate- and lower-resolution cases. The practical role of CryoForge is therefore to automate evidence-supported, bounded and reversible local tasks, reducing routine region-by-region expert work while reserving broad remodeling, cofactor interpretation and conflicting-evidence regions for human assessment.

### Evidence-gated correction balances repair, preservation and escalation

To evaluate CryoForge in recently released complex assemblies beyond the curated benchmark and challenge-derived cases, we examined three cryo-EM structures spanning different initial-model quality and reconstruction-ambiguity regimes: 8X54, the human *γ*-secretase–APP-C99 complex at 2.90 Å; 9BLB, the cagrilintide–CTR–Gs complex at 3.20 Å; and 9W01, the LARS1–IARS1 complex at 3.16 Å [31–33]. Initial models for all three cases were generated using the same initialization workflow used in the main benchmark. CryoForge then operated with the same run-time evidence hierarchy, staged action registry, QC gates and rollback rules. The deposited reference models were unavailable during runtime and were used exclusively for post hoc structural comparison. Whole-model outcomes are shown in Fig. 8, and the corresponding correction footprints, runtime QC changes and unresolved evidence boundaries are summarized in Table 5.

**Table 5.**
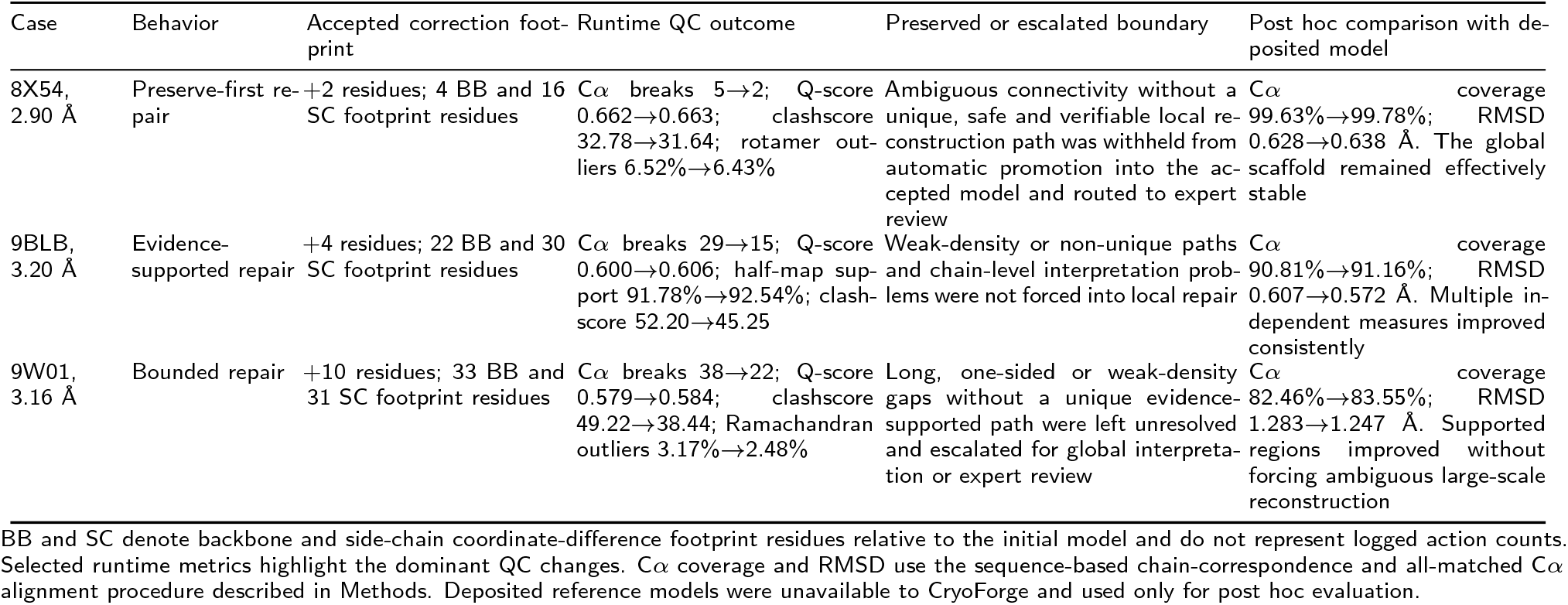
Evidence-gated correction behavior, runtime QC outcomes and post hoc structural comparison across recently released cryo-EM cases. Runtime metrics compare the benchmark-workflow initial model with the final CryoForge post-QC model; deposited-reference metrics were calculated only after processing.

**Figure 8.**
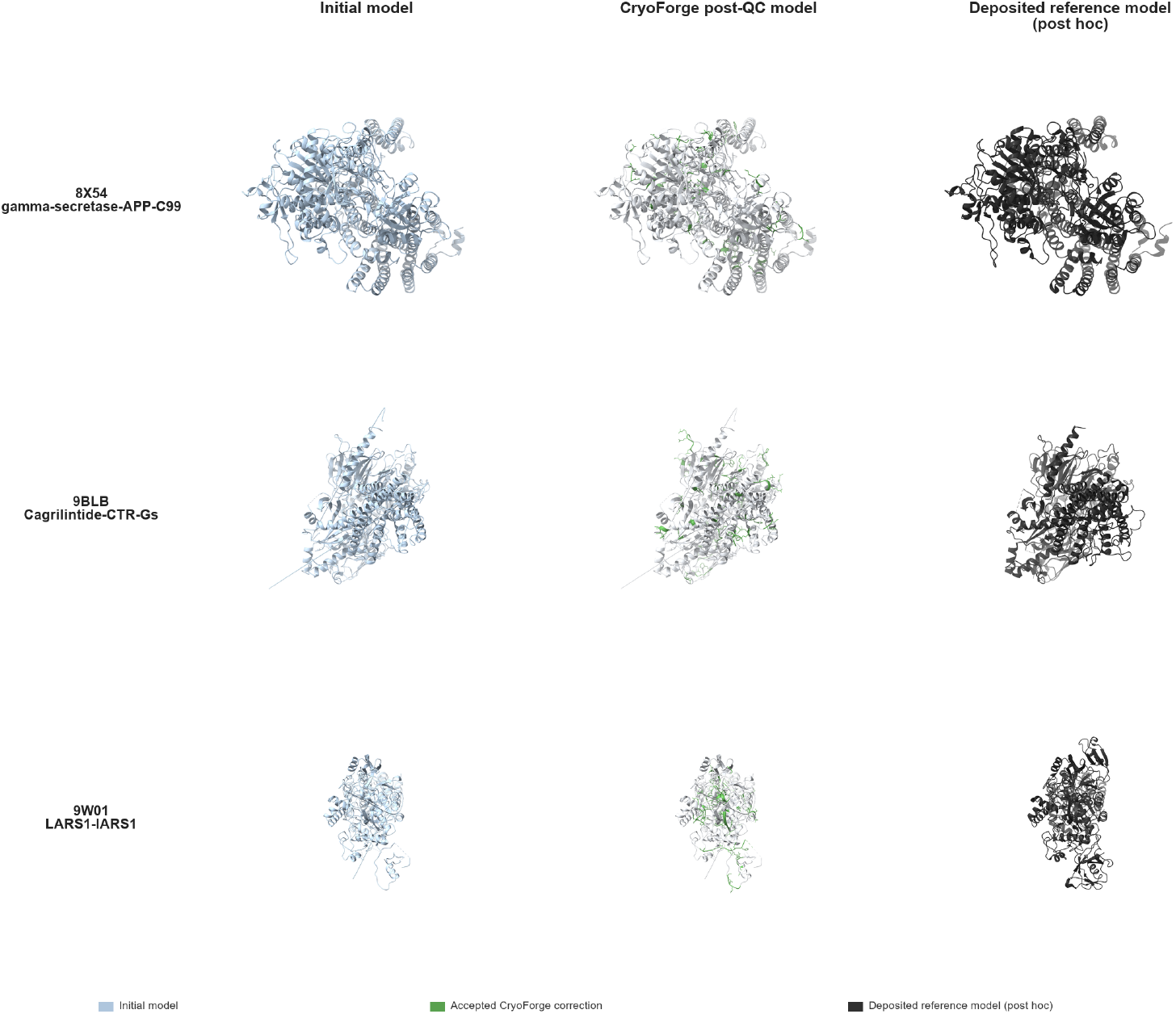
Global structural outcomes of evidence-gated CryoForge correction in three recently released cryo-EM assemblies. Whole-model views are shown for 8X54 (human *γ*-secretase–APP-C99, 2.90 Å), 9BLB (cagrilintide–CTR–Gs, 3.20 Å) and 9W01 (LARS1–IARS1, 3.16 Å). Initial models are light blue. In the post-QC models, retained coordinates are light grey and accepted correction footprints are green. Deposited reference models are dark grey and were used only for post hoc evaluation. Correction scope varied with evidential support; abnormalities lacking a unique, safe and verifiable correction path were withheld from automatic promotion and routed to global interpretation or expert review.

In 8X54, the initial model was already nearly complete and globally well supported, and CryoForge consequently exhibited preserve-first behavior. The accepted footprint comprised two added residues and coordinate differences involving four backbone and 16 side-chain residues, while the number of Cα breaks decreased from five to two. Q-score changed from 0.662 to 0.663, clashscore decreased from 32.78 to 31.64 and rotamer outliers decreased from 6.52% to 6.43%. Post hoc Cα coverage increased from 99.63% to 99.78%, whereas global RMSD remained effectively stable, changing by only 0.010 Å under the all-matched comparison. CryoForge therefore retained the already supported global scaffold rather than introducing broader remodeling. Ambiguous connectivity lacking a unique, safe and verifiable local reconstruction path was withheld from automatic promotion into the accepted model and routed to expert review.

The 9BLB initial model contained more extensive but independently supported local defects, leading to a broader evidence-supported repair footprint. CryoForge added four residues and accepted coordinate-difference footprints involving 22 backbone and 30 side-chain residues, reducing Cα breaks from 29 to 15. Q-score increased from 0.600 to 0.606, half-map support increased from 91.78% to 92.54% and clashscore decreased from 52.20 to 45.25. The same direction of change was observed in the post hoc reference comparison: Cα coverage increased from 90.81% to 91.16% and RMSD decreased from 0.607 to 0.572 Å. Thus, CryoForge accepted a larger correction footprint when density, half-map, connectivity and geometric evidence supported the changes, while weak-density paths, non-unique alternatives and chain-level interpretation problems were not forced into local repair.

The 9W01 initial model had the highest burden of unresolved structural defects among the three assemblies. CryoForge added ten residues and accepted coordinate-difference footprints involving 33 backbone and 31 side-chain residues, reducing Cα breaks from 38 to 22. Q-score increased from 0.579 to 0.584, clashscore decreased from 49.22 to 38.44 and Ramachandran outliers decreased from 3.17% to 2.48%. Post hoc Cα coverage increased from 82.46% to 83.55%, and RMSD decreased from 1.283 to 1.247 Å. CryoForge nevertheless did not attempt to close every remaining gap. Long gaps, one-sided anchors and weak-density regions without a unique evidence-supported reconstruction path were withheld from promotion and routed to Stage4 global interpretation or expert review. The case therefore illustrates bounded repair: the accepted correction scope increased with the amount of repairable evidence but did not cross unresolved evidence boundaries.

Together, the three cases show that CryoForge does not apply a uniform correction scope. It preserved an already well-supported scaffold in 8X54, accepted a broader set of mutually supported corrections in 9BLB and performed substantial but bounded repair in 9W01 while escalating non-unique reconstruction problems. These results indicate that CryoForge treats structural correction as an evidence-constrained decision problem rather than an unconstrained reconstruction task.

## Discussion

This study establishes an evidence-gated post-builder decision layer for cryo-EM structural interpretation. CryoForge reformulates initial-model repair from one-shot coordinate optimization into a region-level process that separates repair proposal from repair acceptance. The framework does not assume that every detectable abnormality should be modified. Instead, it asks which local interpretations are sufficiently supported to change, which should be preserved and which should remain expert-guided. The goal of post-builder correction is therefore not to maximize the number of edits, but to admit only edits that produce evidence-supported improvement. CryoForge provides an acceptance framework for local correction rather than a replacement for comprehensive cryo-EM validation.

The benchmark supports treating initial-model generation and post-builder correction as distinct computational problems. Model generation proposes coordinates from density, sequence and structural priors; correction evaluates whether those existing interpretations remain consistent with local experimental and structural constraints. Residual candidate burden persisted across resolution strata and initialization modes and comprised connectivity, backbone and local-detail defects with different evidence requirements and legal actions. Within this formulation, CryoForge produced substantial local gains while preserving the global scaffold: Stage1 reduced connectivity breaks, Stage2 reduced local geometry defects and most evaluable Stage3 trajectories improved local detail, whereas aligned whole-model Cα RMSD changed little. Stable global agreement does not imply an absence of structural effect, because consequential local errors can be resolved without appreciably changing a whole-model average. The primary endpoint is thus validated resolution of a local target, not global coordinate displacement.

The cross-initialization analysis further shows why this correction layer must be builder-independent. CryoAtom increased model continuity and deposited-reference coverage in the cohort examined here, while shifting the residual burden toward scope, correspondence and local-support assessment. A complete model is not necessarily a fully supported model: every modeled region still requires local experimental evidence [5]. These findings do not imply that one initialization paradigm is intrinsically less reliable, nor should they be generalized to all CryoAtom or AI-derived models. Rather, they show that different builders leave different residual phenotypes and that a useful post-builder system must adapt its triage, legal action space and evidence requirements to the initializer.

CryoForge combines learned prioritization with deterministic evidence acceptance. This division avoids two limiting extremes: fully rule-based control can be less adaptive to heterogeneous structural context, whereas unconstrained learned editing lacks an independent standard for structural acceptance. The learned components narrow the candidate and action search space, but density, geometry, connectivity and neighborhood evidence determine whether a modification is retained. The ablations support this separation: the shared graph representation and learned gate produced most of the reduction in non-improving candidate burden, whereas value learning contributed most when several legal actions required risk-aware selection. The supervisory language model remains restricted to evidence synthesis and expert handoff. Learning therefore assists decision-making without replacing structural validation.

Stopping and rollback are active outcomes of this framework rather than failed attempts at automation. Weak or discontinuous density, long-gap uncertainty, geometry– map tradeoffs, unsupported side-chain completion, sequence/register ambiguity and the absence of a safe legal action define evidence boundaries at which forced repair is unlikely to be reliable. Executability is not equivalent to correctness. By preserving the current model, rejecting a candidate edit or escalating an ambiguous region, CryoForge prevents tool availability from being mistaken for experimental support and keeps unresolved uncertainty explicit. A trustworthy post-builder agent must therefore report not only what it repaired, but also what it protected and why expert judgment remains necessary.

The external-initializer, challenge-derived and recent-complex analyses clarify the practical division of labor between CryoForge and expert-assisted modeling. Across these settings, CryoForge exhibited three complementary behaviors: repair when local evidence supported a bounded change, preserve when an existing interpretation was already reliable and escalate or stop when evidence was insufficient or non-unique. The recent complex-assembly cases made these behaviors explicit across starting models with different residual burdens. Importantly, the near-invariant scaffold in 8X54 and the broader but bounded correction footprints in 9BLB and 9W01 show that correction scope was governed by the amount of locally repairable evidence rather than by a fixed level of algorithmic aggressiveness. Because deposited reference models were unavailable during runtime, the post hoc comparisons assess behavioral calibration rather than reference-guided optimization. The challenge comparisons further showed that CryoForge could reproduce discrete expert-like completeness decisions within a smaller, explicitly recorded coordinate footprint. Expert modelers could integrate global fold continuity, coupled conformational changes, chemical interpretation and cofactor placement across broader regions. The associated metrics also emphasize that geometry optimization, map consistency and deposited-coordinate agreement are related but non-equivalent objectives. HUMAN remodeling often improved clashscore and Ramachandran statistics, whereas CryoForge sometimes retained stronger coordinate-similarity metrics while leaving more residual stereochemical burden. Its conservatism is therefore both a safety property and a limitation: local correction can reduce routine expert workload, but coordinated whole-model refinement may still be required.

Several limitations remain. The benchmark focused on protein-only or protein-dominant single-particle cryo-EM entries within a resolution range suitable for local atomic repair, and claims requiring residue identity, register or side-chain annotation were restricted to sequence-aware settings. Only one contemporary external initializer, three retrospective challenge cases and three recently released complex assemblies were examined. Deposited models are useful post hoc references rather than perfect structural truth, and the challenge archive did not retain exact AUTO-to-HUMAN parent identifiers or chronological expert-operation logs. The current action registry is strongest for local gap closure, short-window backbone correction and side-chain completion; domain-scale retracing, coupled global refinement, ligands, nucleic acids, glycans, heterogeneous conformations and broader chemical interpretation require additional actions and validation data. Future work should test more initializers prospectively, extend the registry, improve uncertainty calibration and integrate structured human feedback for high-risk regions. CryoForge complements rather than replaces de novo builders or expert modelers. More broadly, its results argue that cryo-EM automation should be evaluated not only by how many coordinates are generated, but by whether local interpretations are supported, whether interventions produce defensible net gains and whether uncertainty is handled explicitly. CryoForge thereby moves cryo-EM automation beyond model generation toward evidence-based management of structural interpretation.

## Methods

### Task formulation and CryoForge overview

CryoForge was designed for post-initial-model repair rather than de novo map-to-model generation. Given an initial atomic model, a reconstructed full map, the corresponding half maps, sequence information when available and optional structural priors, the system performs sequential, region-level improvement of the current model. We use closed-loop repair to denote iterative observe–select– execute–validate–update cycles with explicit accept, rollback, stop, data-limited and escalation states, rather than unrestricted autonomous finalization of a structural model.

CryoForge contains two decision components operating under deterministic structural constraints. The GNN/RL component estimates the expected gain and risk of trained repair actions using focal region graphs and off-policy value learning. A supervisory large language model (implemented with Codex/GPT-5.5 in this study) reviews structured evidence, policy scores, QC summaries and action history to summarize complex cases and support expert review [34, 35]. It does not generate coordinate edits or determine final acceptance, which remains subject to the staged QC framework described below.

Candidate edits are evaluated independently before any change is incorporated into the repaired model. This separates learned prioritization and supervisory interpretation from the structural checks required for acceptance.

### Region-level evidence-gated evaluation and acceptance framework

We used a region-level evidence-gated evaluation and acceptance framework to make the repair objective operational. The framework is not intended as a universal validation standard for cryo-EM models. Instead, it specifies how CryoForge decides whether a local post-builder edit is actionable, promotable or unsafe to promote under the available evidence. Its evaluation unit is a candidate residue region rather than a whole model, a single residue or an unconstrained coordinate-editing command. For each region, CryoForge records the local evidence pro-file, repairability state, legal action set, post-action QC requirements and terminal outcome.

Operationally, a region can enter automatic repair only when the defect is localized enough to evaluate and at least one compatible action remains legal after stage, input-readiness, backend and safety masks. A candidate edit is accepted only when required post-action QC supports the intended local improvement without unacceptable regression in map support, half-map consistency, stereochemistry, connectivity or neighboring context. If evidence is insufficient, conflicting, technically unavailable or repeatedly fails QC, the region is assigned to a data-limited, rollback-protected, unchanged or expert-review state rather than being forced into a coordinate change. Thus, stop and rollback decisions are part of the evaluation output, not merely unclassified software failures.

Table 6 summarizes the framework elements used throughout the Methods and Results. The same definitions were used for the paired benchmark, staged repair analysis, learned decision-layer comparisons, non-improving trajectory audit, CryoAtom-initialization compatibility test and recent complex-assembly cases.

**Table 6.**
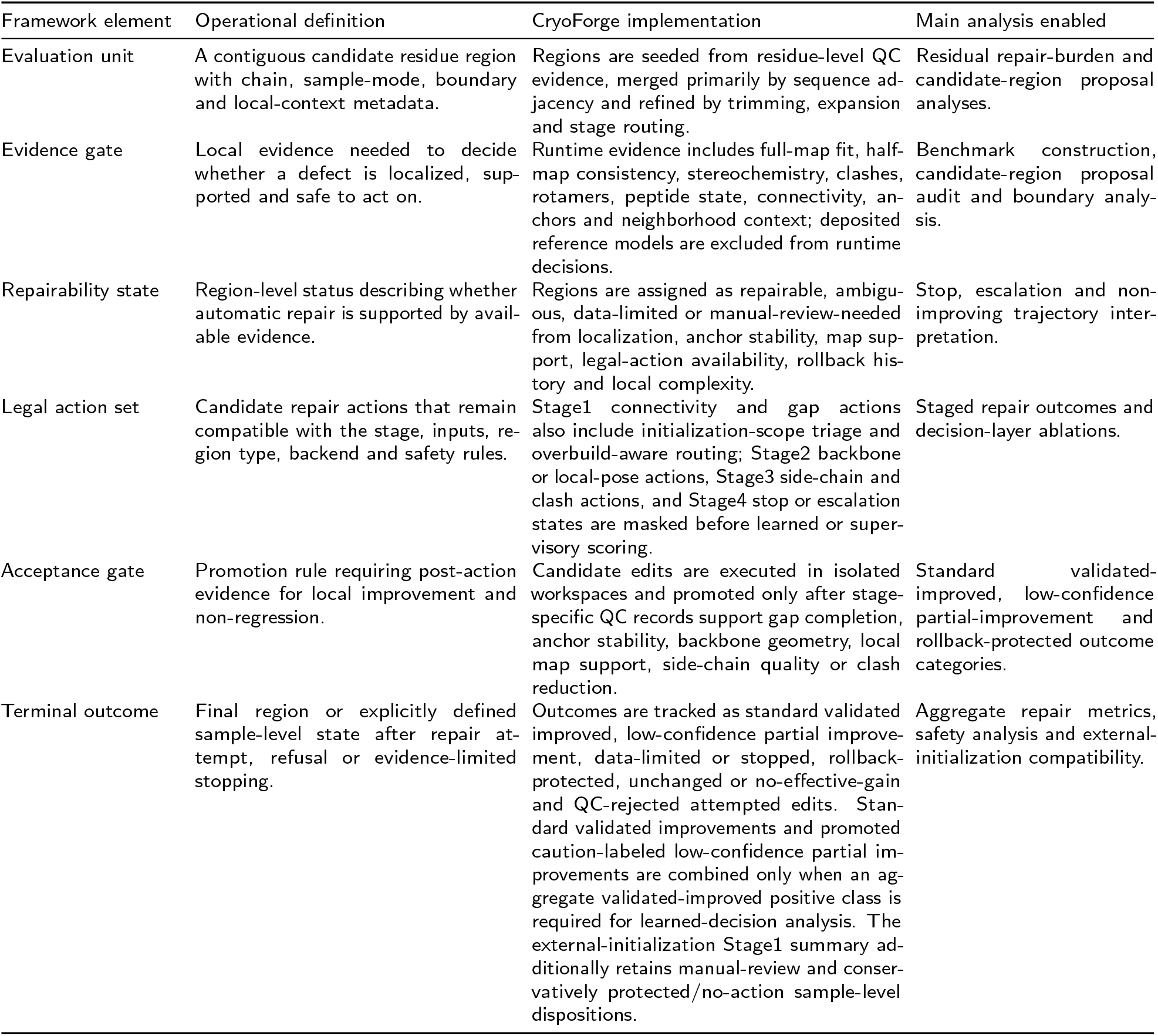
Region-level evidence-gated evaluation and acceptance framework used by CryoForge. The framework defines how local repair candidates are evaluated, which actions are allowed, when candidate edits are promoted and how unsupported cases are recorded.

| Framework element | Operational definition | CryoForge implementation | Main analysis enabled |
| --- | --- | --- | --- |
| Evaluation unit | A contiguous candidate residue region with chain, sample-mode, boundary and local-context metadata. | Regions are seeded from residue-level QC evidence, merged primarily by sequence adjacency and refined by trimming, expansion and stage routing. | Residual repair-burden and candidate-region proposal analyses. |
| Evidence gate | Local evidence needed to decide whether a defect is localized, supported and safe to act on. | Runtime evidence includes full-map fit, half-map consistency, stereochemistry, clashes, rotamers, peptide state, connectivity, anchors and neighborhood context; deposited reference models are excluded from runtime decisions. | Benchmark construction, candidate-region proposal audit and boundary analysis. |
| Repairability state | Region-level status describing whether automatic repair is supported by available evidence. | Regions are assigned as repairable, ambiguous, data-limited or manual-review-needed from localization, anchor stability, map support, legal-action availability, rollback history and local complexity. | Stop, escalation and non-improving trajectory interpretation. |
| Legal action set | Candidate repair actions that remain compatible with the stage, inputs, region type, backend and safety rules. | Stage1 connectivity and gap actions also include initialization-scope triage and overbuild-aware routing; Stage2 backbone or local-pose actions, Stage3 side-chain and clash actions, and Stage4 stop or escalation states are masked before learned or supervisory scoring. | Staged repair outcomes and decision-layer ablations. |
| Acceptance gate | Promotion rule requiring post-action evidence for local improvement and non-regression. | Candidate edits are executed in isolated workspaces and promoted only after stage-specific QC records support gap completion, anchor stability, backbone geometry, local map support, side-chain quality or clash reduction. | Standard validated-improved, low-confidence partial-improvement and rollback-protected outcome categories. |
| Terminal outcome | Final region or explicitly defined sample-level state after repair attempt, refusal or evidence-limited stopping. | Outcomes are tracked as standard validated improved, low-confidence partial improvement, data-limited or stopped, rollback-protected, unchanged or no-effective-gain and QC-rejected attempted edits. Standard validated improvements and promoted caution-labeled low-confidence partial improvements are combined only when an aggregate validated-improved positive class is required for learned-decision analysis. The external-initialization Stage1 summary additionally retains manual-review and conservatively protected/no-action sample-level dispositions. | Aggregate repair metrics, safety analysis and external-initialization compatibility. |

### Benchmark construction and initial-model regimes

Benchmark entries were assembled from publicly deposited cryo-EM map–model pairs with associated full maps, half maps, deposited atomic models and sequence records when available [36–39]. Candidate entries emphasized protein-only or protein-dominant single-particle reconstructions in a resolution range where local atomic repair is meaningful. Runtime decisions used experimental full-map fit, half-map consistency, local stereochemistry, chain connectivity and neighborhood context as primary evidence. AlphaFold-derived or other predicted structures were optional runtime priors rather than ground truth [18], whereas the paired deposited atomic models served as benchmark reference models only for offline label construction, evaluation, compatibility auditing and post hoc analyses.

Initial models were prepared under paired regimes to test whether CryoForge remained useful across different starting-model sources. The full paired benchmark used for initial candidate-region burden analysis contained 849 cryo-EM entries and 1,698 sample-mode records, with one sequence-aware (with-sequence) and one sequence-free (without-sequence or density-dominant) initial model generated for each entry. Entries were balanced across reported resolution buckets, with 283 entries in each of the 2.5–3.0, 3.0–3.5 and 3.5–4.0 Å strata, and each stratum was split into training, validation and test partitions at 70.7%, 14.1% and 15.2% of entries, respectively. This yielded 600 training, 120 validation and 129 test entries; the 600-entry training partition, referred to in Part 2 as the development cohort, corresponded to 1,200 initial-model sample modes across the sequence-aware and sequence-free regimes. The 120-entry validation partition remained separate for model selection and threshold tuning.

A separate external-initialization analysis used CryoAtom-derived initial models as inputs to the frozen CryoForge repair workflow [14]. CryoAtom was selected as a contemporary AI-based builder with a technically distinct initialization route from the benchmark workflow. Benchmark-workflow and CryoAtom-derived initial models were generated or collected for the same 129 held-out test entries and compared by post hoc closeness to the deposited reference model and by initialization-specific stage-entry candidate composition. CryoForge was then applied to the CryoAtom-derived models using the same runtime evidence hierarchy, candidate-region construction, legal-action registry, backend-readiness checks, QC gates and rollback rules. Because this analysis used a single external generator, it was treated as a cross-initialization compatibility test rather than a comprehensive benchmark of current model-building programs.

For the CryoAtom comparison, Cα-level metrics were computed retrospectively after coordinate-frame alignment and chain/residue correspondence mapping using fixed distance criteria listed in Supplementary Methods 1. Cα breaks were normalized per 1000 modeled Cα atoms. Deposited-model Cα coverage used deposited-model Cα atoms as the denominator, whereas matched and unmatched or extra-like modeled Cα fractions used modeled Cα atoms as the denominator. Aligned Cα RMSD and whole-model visualizations used ICP/Kabsch paired coordinates. These metrics were used only to characterize residual-error phenotype or display global alignment and did not guide repair.

For the descriptive phenotype audit in Fig. 6B, classic and CryoAtom candidates were taken from the same 129 held-out entries. Candidate regions were re-evaluated at entry to each stage and assigned to the harmonized phenotype classes shown in the heat maps. Each heat-map column was normalized by the total number of stage-entry candidates for that initialization and stage; the percentages therefore describe within-pool composition and are not estimates of relative repair-method performance or absolute candidate burden. The CryoAtom stage-entry sets contained 8,094 Stage1, 7,288 Stage2 and 400 Stage3 candidate regions.

Stage-entry candidates then passed through readiness, legal-action availability, backend compatibility and high-precision execution-selection gates before any candidate edit was executed. These gates reduced the CryoAtom downstream execution set to six Stage2 and nine Stage3 released region trajectories. Table 3 uses stage-appropriate units: Stage1 reports one sample-level terminal outcome for each of the 129 entries, whereas Stage2 and Stage3 report released region trajectories. Percentages were calculated within each row. Manual review denotes a Stage1 sample-level handoff requiring expert assessment; conservatively protected/no action denotes a sample-level state in which the current coordinates were retained without an unsupported automatic edit. Rollback-protected/QC-rejected denotes an executed candidate edit that was excluded from the promoted model state after post-action QC. Data-limited or stopped, unchanged or no-effective-gain and final-worsened-retained outcomes were zero in these external-initialization rows.

Stage-specific model development used a strict full-QC training subset requiring complete inputs, residue-level quality assessment, candidate-region definitions and stage-entry QC. This subset contained 600 entries and 1,200 paired sample-mode records across the with-sequence and without-sequence regimes. Validation and held-out test records satisfying the same QC requirements were kept separate for model selection, ablation analysis and final performance reporting. Detailed inclusion criteria, split construction and Stage0–Stage4 routing definitions are described in Supplementary Methods 1, and the analysis units and denominators of the principal results are summarized in Supplementary Table 3.

Candidate-region burden was normalized as the number of detected candidate regions per 100 modeled residues in the corresponding model state:

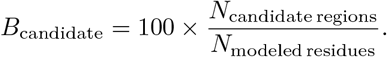

The denominator was the predicted or modeled residue count, pred_residue_count, extracted for each sample-mode, rather than sequence length, deposited-reference residue count, assessed-residue count or QC-passing residue count.

For the resolution-stratified burden summaries, this quantity was first calculated separately for each sample-mode in the 600-entry development cohort. In Fig. 2C, open circles represent the median sample-mode burden within each resolution and initialization group, and thick vertical bars represent the corresponding interquartile range. In Fig. 2D, each heat-map cell reports the macro mean of the sample-mode-level Stage1, Stage2 or Stage3 burden in that group, giving equal weight to each sample-mode rather than pooling candidate-region and residue counts across the group.

Stage-wise major defect-type composition in Table 1 was computed from the 600-entry development cohort’s stage-entry candidate-region sets, not by repeatedly labeling the same initial regions. Stage1 labels were assigned at triage; Stage2 and Stage3 labels were assigned after the preceding repair stage by re-running residue-level QC, rule-based candidate-region proposal and defect typing on the current model. Thus, later-stage labels represent residual, newly exposed or still-actionable local errors. For defect type *d* in stage *s*, composition was defined as

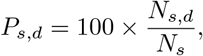

where *N*_*s,d*_ is the number of stage-entry candidate regions assigned to defect type *d*, and *N*_*s*_ is the total number of candidate regions entering stage *s*. The resulting stage-entry sets contained 22,996 Stage1, 9,395 Stage2 and 17,219 Stage3 candidate regions; rare residual side-chain-completion-like cases were grouped with other side-chain defects.

The prespecified frozen proposal audit used the sequence-aware sample modes from the same 600-entry development cohort, subject to prespecified reference-matching eligibility. Deposited-model-derived defect regions were defined from local Cα deviations between the initial model and the paired deposited reference model after chain and residue correspondence mapping in a shared coordinate frame. For each initial-model Cα, the nearest deposited-reference Cα was identified; residues with a nearest-reference distance of at least 3.0 Å were labeled as deposited-model-derived defect residues, and adjacent defect residues were merged into reference defect intervals. Sample-modes with fewer than 50 mappable Cα atoms were excluded from residue-level matching. Rule-proposed candidate regions were audited against these deposited-model-derived regions at region and residue levels. Candidate regions and deposited-model-derived defect regions were represented as residue-index sets:

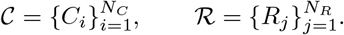

All candidate/reference intersections in this audit were evaluated only for regions mapped to the same chain. Reference-defect region coverage and residue-level recall were defined as

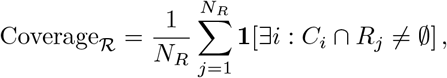

and

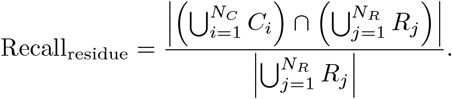

Matched candidate/reference region pairs were assigned by maximum residue-level overlap. For a matched pair *C, R*, intersection-over-union and boundary deviation were defined as

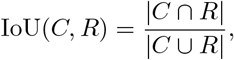

and

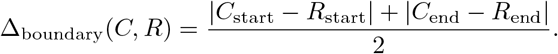

For boundary-accuracy summaries, matched intervals with boundary deviation of at most two residues were counted as boundary matches. The reported reference-defect region coverage, residue-level recall, mean intersection-over-union and median boundary deviation were generated together from the same frozen final audit result and therefore share the same audit eligibility and reference-matching records. These metrics were used only for offline audit of rule-based candidate-region proposal reliability and were not used for runtime repair decisions.

### Recent complex-assembly cases and post hoc model correspondence

The recent complex-assembly analysis used PDB entries 8X54, 9BLB and 9W01 as external case studies [31–33]. Initial models were generated using the same initialization workflow used for the main benchmark, and CryoForge was applied using the same runtime evidence hierarchy, candidate-region construction, staged action registry, QC gates and rollback rules. Deposited reference coordinates were unavailable to the runtime controller and were introduced only after completion of CryoForge processing for global structural visualization, Cα coverage and RMSD calculation. BB and SC counts denote backbone and side-chain coordinate-difference footprint residues between the initial and post-QC models and are not logged action counts.

Escalation was assigned when CryoForge detected a structural abnormality but the available local evidence was insufficient to support a unique, safe and verifiable automatic correction. Such regions were withheld from automatic promotion and routed to Stage4 global interpretation or expert review. Escalation therefore represents an explicit evidence-limited terminal decision rather than an executed repair failure.

For post hoc coordinate comparison, protein-chain sequences from each evaluated model and the deposited reference were globally aligned using scores of 2 for a match, *−*1 for a mismatch, *−*3 for gap opening and *−*0.5 for gap extension. One-to-one chain correspondences were assigned greedily according to the number of aligned identical residues and sequence identity. Chain pairs with sequence identity below 0.25 or fewer than three identical residues were excluded. Cα RMSD was calculated after a single global Kabsch alignment over all sequence-aligned residue pairs for which Cα atoms were present in both models, without trimming, outlier rejection or local selection. Deposited-model Cα coverage used sequence-mapped deposited Cα positions as the denominator. Coverage and RMSD were recomputed independently for the initial and post-QC models under the same correspondence procedure; newly added residues could therefore enter the post-QC matched set. These metrics were interpreted jointly as post hoc global-agreement measures rather than as fixed-atom paired displacements, and neither metric was used for runtime repair decisions.

### Candidate-region construction from residue-level evidence

Each repair round began with standardized residue-level validation of the current model. CryoForge combined local density-fit, half-map consistency, stereochemical, clash, rotamer, Ramachandran, peptide, correspondence, connectivity and neighborhood-context features into a residue-level evidence table. Candidate regions were seeded from low-quality or structurally suspicious residues within the same sample, initial-model mode and chain, then merged primarily by sequence adjacency. Clean edge residues could be trimmed and nearby suspicious residues could expand the boundary during GNN refinement, whereas gap state and anchor suitability were assessed during Stage1 routing.

Region labels summarized both dominant defect mechanism and repairability. Dominant types included backbone-tracing-like, register-assignment-like, side-chain-packing-like, weak-density-loop-like and mixed regions, with connectivity, backbone and register defects prioritized before side-chain repair. Repairability was represented as repairable, ambiguous, data-limited or manual-review-needed according to localization, anchor stability, map and half-map support, legal-action availability, repeated roll-back and complex local context. Candidate regions were managed through a priority queue refreshed after post-action QC; the complete feature dictionary, thresholds, region-merging rules and subtype definitions are provided in Supplementary Methods 2.

For external-initialization triage, Stage1 region labels were further collapsed into phenotype-level classes used in the CryoAtom analysis. High-confidence local gap routes required short, anchored intervals with usable map and half-map support and no post-action connectivity or geometry regression. Low-confidence or data-limited gaps were short and modelable but had weak map support or relied primarily on sequence or runtime-reference anchors; overlong gaps were routed to Stage4, expert review or higher-level rebuild rather than forced through local completion. Weak-density and very-low-support calls combined Q-score, FSC-Q and full/half-map evidence, whereas reference-guided low-confidence repair required sufficient runtime reference–target Cα correspondence and local map compatibility. Runtime reference conflict blocked reference-guided actions but did not by itself make a region unrecoverable. Partial-connectivity, overbuilt-like segment and scope-expanded labels captured local continuity defects, low-support modeled fragments, high break-density fragments or many short low-support chains, and were treated as audit/triage phenotypes rather than completion targets. Runtime scope filtering or deletion was conservative and applied only after chain-level quarantine and fraction-of-model safeguards. Detailed cutoffs are provided in Supplementary Methods 2; deposited-reference information was used only for offline benchmark evaluation, training audit or visualization and not for runtime routing.

### Staged action registry and QC-gated execution

Repair was exposed as a staged environment rather than a single unrestricted editing step. Stage0 performed sample-mode triage and assigned each record to no action likely good, Stage1 connectivity or gap repair, candidate-region proposal audit or data-limited handling. Stage1 targeted connectivity defects, residue-register inconsistencies and chain-continuity problems using GapFill routing with anchor, gap-length, map-support and backend-readiness checks. Stage2 focused on backbone placement and local-pose repair under map and geometry constraints, Stage3 handled side-chain placement, clash resolution and rotamer correction, and Stage4 represented data-limited or ambiguous regions requiring masking or expert review.

Final evidence and reports were assembled after staged repair.

Actions were selected from a registry rather than generated as free-form commands. The mature registry covered Stage1 sequential gap filling, Stage2 backbone cleanup, anchor-locked idealization and local real-space refinement, and Stage3 rotamer repair, clash-aware repacking, side-chain regularization and sequence- or runtime-reference-guided completion. Every candidate action was checked for legality, backend availability, required inputs, region compatibility, recent failure history and user permissions before scoring; illegal or unavailable actions were masked for learned, rule-based and supervisory components. Complete action names, backend mappings, extension schemas and transition tables are provided in Supplementary Methods 3.

All trained and extension actions used the same legality and post-action QC rules. Promotion required deterministic legality filtering and stage-specific QC: gap completion and anchor stability in Stage1, backbone geometry and local non-regression in Stage2, and side-chain completeness, rotamer, clash and density support in Stage3. Stage-level outcomes were classified as improved, partial or low-confidence improved, rollback, unresolved mask or action-inapplicable. Rollback or escalation was required when QC regressed, non-local damage was introduced, geometry or connectivity worsened, the output was invalid, required inputs were unavailable or evidence remained insufficient for safe promotion.

A region was released from Stage1 to Stage2 or Stage3 only when it was localized, not quarantined, had all required inputs and backend readiness, retained at least one legal action and had sufficient map, anchor and neighborhood evidence for stage-specific QC. Regions failing these checks were reported as data-limited, masked or manual-review rather than being forced into local refinement. Low-confidence partial improvement was assigned when a Stage3 candidate completed successfully and improved side-chain completeness, geometry, clash or rotamer evidence while preserving backbone stability, neighboring-region stability and local map or half-map support, but retained density, rotamer or local-resolution ambiguity. Prespecified regression, improvement, Q/FSC and runtime-reference-fit tolerances are reported in Supplementary Methods 3. Under weak map evidence, Stage3 completion could be labeled low-confidence or geometry-completion improvement, but not high-confidence map-resolved side-chain repair.

### GNN/RL value learning for trained repair actions

For trained registry entries, CryoForge used a GNN/RL value model to prioritize executable local repair actions. The policy state combined a focal region graph, region- and sample-level context, the legal action mask and dominant error-type features. Graph nodes and edges encoded local density-fit, half-map, geometry, connectivity, prior-compatibility and repair-history descriptors, and a graph neural network encoder produced region embeddings used to estimate action value, expected gain and expected risk [40].

Learning was stage-specific. Stage0 predicted triage routes from sample-mode and aggregated region features. Stage2 and Stage3 used a GNN region head for actionability filtering followed by a contextual Q head over legal actions, whereas Stage1 GapFill routing remained dominated by deterministic anchor and gap-length constraints. Training used supervised warm-starting followed by replay-based value refinement with target-network stabilization and Double-DQN-style action evaluation [41, 42]. Replay targets encoded post-QC outcome utility using ordinal rewards for partial improvement, validated or map-supported improvement, rollback or worse outcomes, and preflight-blocked or unexecuted actions. At inference time, the learned policy scored only legal trained-registry actions; untrained extensions were routed through deterministic compatibility rules, user request or supervisory arbitration under the same QC gate. Architecture, replay-buffer construction, reward normalization and ablations are described in Supplementary Methods 4.

### Supervisory LLM and evidence review

The supervisory LLM was implemented with Codex/GPT-5.5 in this study and operated over structured evidence rather than raw coordinates. Its inputs included region descriptions, available repair choices, GNN/RL scores, QC summaries, previous outcomes and user constraints; its outputs were evidence summaries, recommendations for complex cases and suggestions for manual review.

The supervisory component handled high-risk or conflicting-evidence cases but could not introduce unavailable repairs, bypass required inputs, override hard QC failures or accept an unvalidated model state. A dedicated CryoForge skill layer provided repair descriptions, QC checklists, reversal requirements and reporting templates. Skill instructions and prompt-level safeguards are provided in Supplementary Methods 4.

### Decision baselines and evaluation

CryoForge was evaluated on held-out full-QC sequence-aware cryo-EM map–model entries stratified by reported resolution, dominant defect class, repairability and action type. The main comparisons included the unmodified initial model, a rule-only staged repair strategy, GNN-only and GNN/RL decision strategies, and the full CryoForge system.

The GNN-only comparator predicted the stage/error route and repairability labels, used the learned graph score for region filtering and actionability assessment, and ranked legal actions by predicted immediate gain. After legality masking, GNN-only selected the action with the highest predicted gain. The GNN/RL selector instead ranked the same legal actions by predicted Q, incorporating expected gain, expected risk and downstream utility. Prespecified stability and high-risk checks allowed either learned selector to skip marginal or unsafe edits. This separation enabled comparison of supervised immediate-gain ranking with risk-aware value-based action ranking.

For learned-decision analyses, all strategies used the same candidate-region pool, legal-action registry, QC gates and rollback rules. Region filtering was assessed by precision, recall, average precision and non-improving-candidate retention. The aggregate validated-improved class, including promoted low-confidence partial-improvement subtypes that passed all non-regression safeguards, was treated as the positive repair-target class. A non-improving candidate was operationally defined as a candidate that did not enter this aggregate class after QC. For a score threshold *τ*, let *S*_*τ*_ denote retained candidate regions and let *P* denote candidate regions in the aggregate validated-improved class. Region-filtering precision and recall were defined as

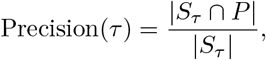

and

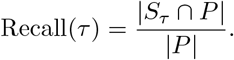

Average precision was computed from the threshold-swept precision–recall curve as

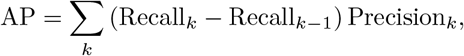

with thresholds ordered by decreasing candidate score. Non-improving-candidate retention at threshold *τ* was defined as

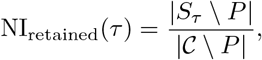

where *c* is the full rule-generated candidate-region pool. Action selection was assessed by match to the post hoc oracle-best legal action and harmful-action rate. For candidate regions with multiple legal actions, action-match rate was defined as

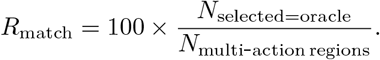

The oracle-best action was computed from observed post-QC outcomes and used only as an upper-bound reference,not as a runtime-available policy. Harmful-action rate was defined as

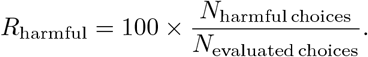

These analyses focused on Stage2 and Stage3, where .candidate filtering and multi-action selection were most informative; Stage1 gap and connectivity routing remained dominated by deterministic readiness, anchor and gap-length constraints.

A released repair-region trajectory was a region-level record that passed stage-entry readiness and entered QC-gated repair handling, including legal action execution, rollback-protected rejection, data-limited stopping or unchanged/no-effective-gain termination. Initial no-action likely good records and regions screened out before stage-entry readiness were not counted in the released-trajectory denominator.

For terminal-outcome reporting, standard validated improvements and caution-labeled low-confidence partial improvements were displayed separately. Standard validated improvement denoted a promoted QC-supported repair without the residual ambiguity required for the low-confidence label. Low-confidence partial improvement denoted a promoted repair that passed all non-regression safeguards but retained explicitly recorded density, rotamer or local-resolution ambiguity. The two promoted classes contained 22,191 and 970 trajectories, respectively, and were combined into the aggregate validated-improved class only for learned-decision positive labels and summaries requiring the aggregate definition. Other stage-level transition labels were collapsed into data-limited or stopped, rollback-protected or QC-rejected, unchanged or no-effective-gain and final worsened retained. Rollback-protected and QC-rejected attempted edits were combined because both ended with the attempted edit excluded from the promoted model state. For final outcome class *o*, outcome proportion was defined as

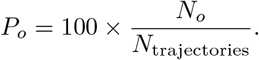

Representative staged examples were selected post hoc from standard validated-improved cases with complete initial, repaired, deposited-reference, map and stage-evidence files. Cases were prioritized for clear visual interpretability, complete Stage1/2/3 evidence and representative stage-specific repair behavior. They were not randomly sampled and were not used to estimate aggregate performance statistics. For a defect class *q*, before–after defect reduction was reported as

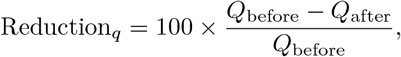

where *Q* denotes the stage-specific defect count or normalized defect rate. Stage1 connectivity breaks per gap and Stage2 geometry defects per region were computed as

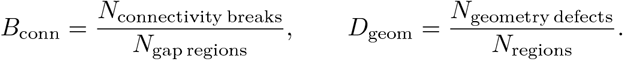

The Stage3 local-detail-gain rate shown in Fig. 3C was defined as

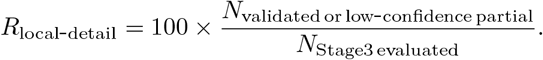

The numerator included standard validated improvements and caution-labeled low-confidence partial improvements under the stage-specific density, geometry, clash, rotamer and non-regression criteria. Global backbone preservation was summarized by aligned Cα RMSD to the deposited reference model,

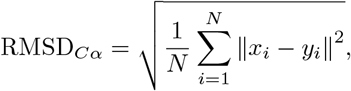

where *x*_*i*_ and *y*_*i*_ are aligned Cα coordinates in the repaired/current model and deposited reference model, respectively. The reported paired change was the mean across models of RMSD_*C*α,post-QC_ *−* RMSD_*C*α,initial_.

For stage-specific evidence-boundary analysis, each event was assigned to the structural factor that most directly prevented a validated improvement. Table 2 included 834 Stage1 events, 1,819 Stage2 non-improving events and 1,249 Stage3 non-improving events. Categories were long gap or large missing segment, weak or discontinuous density, geometry–map tradeoff, absence of a safe repair, insufficient structural support and sequence/register ambiguity. When several signals were present, the dominant category followed a prespecified order based on input validity, availability of a plausible repair, map support, gap length, connectivity or register uncertainty, backbone defects and side-chain defects. Deposited-model comparisons were used only retrospectively. Weak map evidence did not automatically preclude repair: short, anchored and modelable regions could still proceed when the remaining evidence supported a cautious correction.

Local density support across profiled gaps was quantified using an anchor-normalized density proxy. For each profiled residue, the candidate local-path Cα coordinate was used to sample the full-map density value at the nearest map voxel. The mean density over the two flanking anchor segments was used as the 100% reference level, and residue-level relative support was defined as

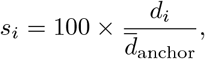

where *d*_*i*_ is the full-map density at residue *i* and *d*_anchor_ is the mean anchor density. Gap-level support was the mean relative support across residues inside the gap:

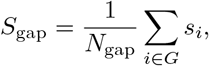

where *G* is the set of profiled gap residues and *N*_gap_ =| *G* |. For the representative boundary analysis, 50% of anchor-normalized density was used as the low-support cutoff. A gap was considered weakly supported when most gap residues fell below this cutoff while the flanking anchors retained substantially higher support.

Deposited-model coverage was quantified retrospectively by recording whether each position in the profiled interval had a corresponding modeled residue in the deposited model. The supported fraction for an interval *I* was defined as

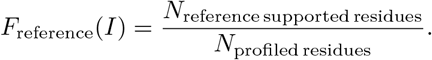

Geometry–map tradeoff was assigned when an attempted edit reduced local geometry-defect counts but decreased local map support beyond the acceptance tolerance. QC-rejected attempted edits were completed candidate edits that failed the post-action safety gate and were not promoted; the protected pre-action model was retained by rollback. Boundary composition was calculated within each stage-specific boundary/non-improving event class as

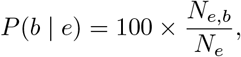

where *N*_*e,b*_ is the number of events in stage-specific event set *e* assigned to boundary reason *b*, and *N*_*e*_ is the total number of events in that set. Thus, boundary percentages describe composition within each Stage1 triage-boundary, Stage2 non-improving or Stage3 non-improving event set rather than the global frequency of each boundary type across all repair trajectories. Candidate regions were reevaluated at stage entry, so these event counts are not additive across stages and remain separate from the 26,153-trajectory denominator used for aggregate repair outcomes.

Evaluation metrics were grouped into structural, decision and safety categories. Structural metrics included local density-fit improvement, half-map consistency, geometry improvement, severe-error reduction and region-resolution rate. Decision metrics included action-selection accuracy, value-prediction error, realized reward, gain/risk calibration and selection stability. Safety metrics included harmful-action rate, reversal rate, false-accept rate and the frequency of unresolved high-risk regions.

Core metrics were defined at the region, action or report level, and deposited reference models were not used for runtime action choice. The best action for legal-action top-1 accuracy was defined by the realized deployable post-QC outcome in exhaustive or replayed outcomes. Harmful actions were edits that increased stage-local severity, regressed map or half-map support, worsened geometry, clash, peptide or connectivity evidence, introduced non-local damage or produced invalid output. Deposited-model-derived candidate-region proposal audits were restricted to post hoc defect enrichment and within-candidate reference-overlap comparisons. Severity bins were defined from the composite severity score *s* as

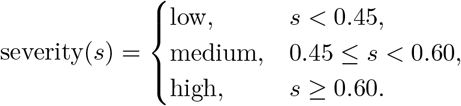

Statistical analyses, seed handling, software versions, replay-buffer snapshots, checkpoints and output schemas are provided in Supplementary Methods 5.

## AI Tool Use Declaration

CryoForge includes a supervisory LLM, implemented with Codex/GPT-5.5 in this study, as part of the research system described in the Methods. This component reviewed structured evidence, available repair choices and QC summaries; it did not directly edit atomic coordinates, introduce unavailable repairs or override deterministic QC failures. Separately, large language model tools were used during manuscript preparation only for LaTeX organization and consistency checking. All scientific claims, analyses, figures, data interpretation and final text were reviewed, verified and approved by the human authors.

## Data Availability

The cryo-EM maps, half maps where available, deposited atomic models and associated sequence records analyzed in this study were downloaded from public structural biology repositories, including the Electron Microscopy Data Bank (EMDB) and the Protein Data Bank (PDB) [37–39]. Accession codes and benchmark split manifests are provided through the public CryoForge project repository. Source data underlying the figures and quantitative results are provided with this paper.

## Code Availability

The CryoForge source code, analysis scripts, figure-generation scripts, benchmark-manifest schemas and example configuration files are available Additional implementation details including benchmark construction, candidate-region definition, action-registry design, GNN/RL learning, LLM supervision and reproducibility records, are provided in Sup plementary Methods 1–5.

## Author Contributions

W.F. and R.H. conceived the study. W.F. developed the CryoForge framework, assembled the benchmark, performed the computational analyses, prepared the figures and drafted the manuscript. R.H. supervised the study, contributed to method design and interpretation, and revised the manuscript. All authors reviewed and approved the final manuscript.

