## Supplementary Information for "CryoForge: A Self-Correcting Agent for Cryo-EM Model Building That Learns When to Act and When to Stop"

### Supplementary Methods 1. Benchmark construction, data splits and initialization modes

#### Benchmark design

CryoForge was evaluated using a resolution-stratified benchmark designed to measure local repair burden after cryo-EM initial-model generation while preventing entry-level information leakage. The benchmark contained 849 protein-only or protein-dominant single-particle cryo-EM entries with reported global resolutions between 2.5 and 4.0 Å. Entries were distributed equally across three resolution intervals: 2.5–3.0, 3.0–3.5 and 3.5–4.0 Å, with 283 entries in each interval. The cryo-EM entry, rather than an individual initial model, region or repair attempt, was used as the independent split unit.

Each entry was assigned to one of three fixed partitions: 600 entries for training and method development, 120 for validation and 129 for held-out testing. Within each resolution interval, the corresponding allocation was 200 training, 40 validation and 43 held-out test entries. All derivative records from an entry, including initialization modes, residue evidence, candidate regions, action attempts, replay records and post hoc deposited-gold annotations, inherited the partition of the parent entry. Consequently, different initialization modes from the same cryo-EM entry could not occur in different partitions.

#### Inclusion and exclusion criteria

Entries were eligible when the full cryo-EM map, deposited atomic model, sequence metadata and map/model identifiers could be paired unambiguously and read by the standardized preprocessing workflow. Protein-dominant complexes were retained when the protein component represented the intended repair target. Entries were excluded when the reported resolution was outside the prespecified range, the full map or deposited model was unreadable, map and model identifiers could not be reconciled, the coordinate frame could not be resolved, or the target was outside the protein-focused scope of the current agent.

Half maps, Q-score, FSC-Q and other local QC products were treated as evidence components rather than silently imputed requirements. Their availability and reliability were recorded explicitly. A missing local metric therefore represented unavailable evidence, not an automatically failed structure. Analyses requiring a particular metric used an eligible subset defined before calculation and reported the corresponding denominator. This distinction was important because repairability can remain evaluable from geometry, connectivity and full-map evidence even when one local metric is unavailable, whereas claims about half-map consistency require usable half-map-derived evidence.

#### Sequence-aware and sequence-free initialization

Each benchmark entry was represented by two initial-model modes. In the sequence-aware mode, the target sequence was supplied during initial-model generation. In the sequence-free mode, the same map was modeled without supplying the target sequence to the initialization procedure. The two modes therefore formed a paired contrast within each entry rather than two independent datasets. Across the complete benchmark, this produced 849 sequence-aware and 849 sequence-free sample-modes, for a total of 1,698 initial-model structures. The training, validation and held-out partitions contained 1,200, 240 and 258 sample-modes, respectively.

The principal initial-model workflow used ModelAngelo v1.0.16 (commit ffb872cb) with downstream PHENIX processing (PHENIX 2.0-5936-cuda12). Sequence availability at model generation and sequence availability to a later registered repair action were recorded separately. A sequence-free initial model could therefore be evaluated as sequence-free while still permitting a later sequence-dependent action only when sequence use was explicitly allowed by the action registry and sample metadata. The complete commit identifier is retained in the frozen software manifest.

The 600-entry training partition was used for detector development, action-registry maturation, replay-buffer construction and learned-model fitting. The 120-entry validation partition was used for checkpoint selection, threshold calibration and policy freezing. The 129-entry held-out partition was not used for training, reward tuning, threshold adjustment or registry expansion. Analyses that used only one initialization mode or a subset of metric-compatible entries state this restriction explicitly. The pooled full-QC repair-trajectory corpus used to describe action and QC outcomes in Fig. 3 is a development-scale trajectory analysis and is not presented as a held-out generalization estimate. Held-out performance and external-initialization analyses use the frozen 129-entry test partition.

#### External-initialization compatibility analysis

To test whether CryoForge could operate beyond the principal ModelAngelo–PHENIX initialization workflow, CryoAtom-derived initial models were generated for the same 129 held-out entries and entered through the common compatibility and stage-routing procedures. CryoAtom was treated as an external source of initial coordinates, not as a separate repair branch with a different definition of success.

Before local repair, each external model was checked for map-coordinate compatibility, chain and residue correspondence where available, Cα break burden, modeled Cα coverage, unmatched modeled content and evidence of overbuilt-like or scope-expanded segments. Alignment metrics, including iterative closest point or Kabsch-aligned Cα RMSD, were computed only for compatible matched subsets and were used for post hoc evaluation. They were not used to drive runtime repair.

The compatibility audit distinguished local defects from problems outside the scope of local repair. Coordinate-frame failures, extensive domain displacement, chain/copy correspondence ambiguity and large unsupported scope expansion were routed to global review or manual assessment. Local gaps, backbone defects and side-chain defects could proceed to the common Stage1–Stage3 framework if they passed the corresponding repairability and legality checks. Weakly supported modeled fragments could be quarantined or conservatively removed only when the evidence and post-action QC safeguards required for that action were satisfied.

#### Runtime evidence and deposited-gold audit

The deposited atomic model associated with each benchmark entry is termed the deposited gold model in this study. It was used only for offline label construction within the permitted training partition, post hoc benchmark audit and manuscript evaluation. It was not available to the runtime Stage0–Stage4 controller, legality masks, GNN or GNN/RL inference, coordinate-generation backends, QC promotion decisions or supervisory language model.

Runtime decisions used deployable evidence: the full map, half-map-derived evidence when available, Q-score and FSC-Q when available and reliable, stereochemistry, backbone connectivity, local neighborhood context, sequence when legally available, action history and a permitted structural prior only when that prior was locally compatible with the map. All source-data fields derived from the deposited gold model were labeled as post hoc. This information boundary was enforced both in the software interfaces and in the interpretation of results.

### Supplementary Methods 2. Residue-level evidence, candidate-region construction and repairability labels

#### Evidence representation

CryoForge first converts a model and its associated maps into residue-level evidence. The evidence representation includes residue identity and sequence position; local full-map support; Q-score; FSC-Q and half-map consistency; backbone and peptide geometry; rotamer, clash and atom-completeness flags; peptide C–N and adjacent Cα–Cα connectivity; fragment boundaries; neighboring suspicious residues; unresolved masks; previous accepted or rolled-back edits; and compatibility with a permitted prior when such a prior is available and locally supported by the map.

Each metric is accompanied by availability and reliability flags. Missing Q-score or FSC-Q is therefore not encoded as a poor score. Instead, the region carries incomplete-evidence status, and the repairability gate determines whether the remaining evidence is sufficient for a restricted action or requires data-limited routing. This prevents technical missingness from being converted into a structural negative label.

Deposited-gold distances and interval overlaps were added only to offline audit tables. These fields were excluded from runtime feature matrices, action masks and promotion decisions.

#### Suspicious-residue seeds and region construction

Residues were seeded as suspicious when one or more stage-relevant signals exceeded the frozen policy threshold. Signals included chain breaks, implausible peptide links, local geometry outliers, short torsion anomalies, missing or incomplete side chains, clashes, poor rotamers, low local map support, low Q-score, low FSC-Q, weak half-map agreement, fragment boundaries, unbuilt-density signals and abnormal local neighborhood context. Low density alone was not treated as proof of model error. A residue with weak density but acceptable connectivity and geometry could be labeled data-limited or stable rather than repairable.

Suspicious residues were merged only within the same chain. Adjacent residues were joined directly, and selected residues separated by no more than two positions in sequence order could be merged when they shared a continuous evidence pattern. A one-residue interruption could therefore be bridged within a local event, but regions were not joined across chains unless the sample was being evaluated for an explicit chain/copy correspondence or global-scope problem.

Initial region boundaries were refined by three operations. First, clean terminal residues that did not contribute evidence or anchor context were trimmed. Second, nearby residues were added when needed to represent gap anchors, peptide continuity, local torsion context or side-chain packing. Third, the graph-based boundary model could retain, split, expand or suppress a candidate using deployable residue and neighborhood features. An unresolved upstream region produced a mask that prevented later-stage actions inside the affected interval, with a local buffer when required to protect neighboring geometry. Independent regions outside the mask remained eligible for downstream detection.

#### Detection is distinct from repairability

A candidate region denotes evidence of a possible local defect; it does not imply that a safe automated repair exists. CryoForge therefore applies a repairability gate after detection. The gate assigns one of the following states:

1. **Repairable, map-supported:** the region has an interpretable target, stable local context and sufficient map or half-map evidence for a legal action.
2. **Repairable with locally compatible prior support:** map evidence is weak or incomplete, but anchors are stable and a permitted prior agrees locally with the map and surrounding model. Any accepted edit is labeled low-confidence or prior-guided.
3. **Ambiguous:** the evidence supports a defect signal but not a unique boundary, stage or repair path.
4. **Data-limited:** density is weak or discontinuous, half-map evidence is insufficient, no unique path is identifiable, or the region is likely flexible or disordered.
5. **Manual review or higher-level recovery:** the problem involves global placement, domain orientation, chain/copy correspondence, extensive scope mismatch, an overlong gap, repeated rollback or an unavailable backend.

This separation is central to the workflow. It allows CryoForge to detect a biologically meaningful weak-density or missing-fragment region without forcing an edit that the data cannot support.

#### Frozen structural and map thresholds

The frozen detector used Q-score ≥0.40 as a residue-level supported-density indicator. Mean FSC-Q <0.25 or minimum FSC-Q <0.10 triggered low-density review when FSC-Q was available and reliable. The frozen FSC-Q discrepancy-support flag additionally used |FSC-Q| ≤0.50 according to the recorded QC policy. Because the numerical meaning of FSC-Q depends on the implementation and preprocessing route, all analyses used the policy version stored with the source table rather than mixing values from different calculators.

A peptide C–N distance >2.00 Å was treated as a hard break signal. The expected adjacent C–N interval was 1.15–1.55 Å. An adjacent Cα–Cα distance >5.00 Å was treated as a hard Cα break signal, whereas 3.40–4.20 Å was considered the expected local interval. Peptide ω values with 35° < |ω| <145° were flagged as suspicious. Non-neighbor heavy-atom distances <1.60 Å generated severe clash evidence.

For gap repair, routine high-confidence map-first filling was limited to gaps of 1–8 missing residues with stable anchors and usable local map evidence. Gaps of 9–12 residues could enter conditional prior-guided or low-confidence repair only when anchor geometry, local prior compatibility and backend readiness were sufficient. Gaps >12 residues were outside the mature automatic local gap-filling scope and were routed to Stage4, global recovery or expert review. A separate map-first unbuilt-density tracing route could consider a local unbuilt-density span up to 24 residues, but this was a routing ceiling rather than an assurance that a continuous gap would be filled.

A permitted prior was considered locally usable when at least three local Cα pairs could be matched and target coverage was ≥0.30. The stricter Stage0 prior-support condition required successful adaptation and either local coverage ≥0.45 or at least four matched Cα pairs. A prior that conflicted with the map was marked unusable for that region and could not determine the repair or its acceptance.

#### Threshold derivation and freezing

Frozen thresholds belonged to three provenance classes. Peptide-link distances, adjacent Cα distances, peptide ω ranges and severe-clash distances were fixed structural safety rules derived from established stereochemical expectations. Gap-length ceilings, candidate-merging distance and local-prior coverage requirements were development rules established on the 600-entry training partition while maturing the corresponding selectors and action backends. Learned route, actionability and operating-point thresholds were calibrated on the 120-entry validation partition after model fitting. No threshold was selected or changed using the 129-entry held-out test partition. The threshold class and QC-policy version are retained in the source manifest.

#### Defect taxonomy and stage assignment

Candidate regions received an expert-readable primary class and an action-oriented subtype. Primary classes included missing/unbuilt-fragment or connectivity defects, backbone-tracing-like defects, register or chain-correspondence ambiguity, backbone/local-pose defects, side-chain or local-clash defects, weak-density/data-limited regions, overbuilt-like or scope-expanded segments and mixed/ambiguous regions.

Stage1 subtypes included false-link-like connectivity, high-confidence local gap, low-confidence data-limited gap, partial connectivity, missing/unbuilt density, overlong gap, register ambiguity, fragment stitching and overbuilt-like scope. Stage2 subtypes included short torsion window, peptide-flip-like geometry, local backbone pose, gap-insert backbone pose and local geometry regularization. Stage3 subtypes included side-chain rotamer, side-chain clash/pose, side-chain atom completion and gap-insert side-chain completion.

Safety and upstream dependency determined the label hierarchy. Input or coordinate-frame failure took precedence over local repair. Unresolved Stage1 connectivity or register ambiguity took precedence over Stage2/Stage3 labels within the same interval. Data-limited evidence and absence of a unique path could stop an action at any stage. Stage2 was considered only after connectivity was adequate or the Stage1 region was masked; Stage3 was considered only after the local backbone was sufficiently stable.

#### Stage0 routing criteria

Stage0 is a sample-level triage step rather than a quality score. A sample could be routed to no action, Stage1, Stage2, Stage3, data-limited management, selector audit or global recovery. The frozen no-action-likely-good route required no released candidate region, Q-score $\geq 0.55$, FSC-Q $\geq 0.20$, no detected chain break and a severe-residue fraction $<0.05$. If no region was released but these QC conditions were not all satisfied, the sample entered selector audit rather than no action. The data-limited route required the number of data-limited regions to be at least max(1, floor(0.75 × total region count)), together with Q-score $<0.45$ and FSC-Q $<0.08$. The global-recovery route required all five conditions: Q-score $<0.18$, FSC-Q $<0.02$, severe-residue fraction $>0.70$, $>50$ candidate regions and maximum candidate length $>80$ residues. Samples with detected regions but no action-ready region entered selector/repairability audit.

Samples entered Stage1 when a Stage1 region was present, a chain break was detected or a candidate region extended over at least 25 residues and represented an upstream interpretation problem. These sample-level thresholds did not override region-level repairability. A sample routed to Stage1 could still contain regions that were quarantined, sent to Stage4 or excluded from automatic repair.

#### Post hoc candidate-region audit

Candidate-region proposal quality was assessed against deposited-gold-derived defect intervals in an offline audit. A modeled Cα atom with nearest deposited-gold Cα distance ≥3.0 Å was used as a post hoc defect seed after coordinate-compatible matching. Samples required at least 50 mappable Cα atoms for this audit. Deposited-gold-derived defect residues were merged into intervals allowing a one-residue interruption.

For each deposited-gold-derived interval, the candidate region on the same chain with maximum sequence-interval intersection-over-union (IoU) was selected. Unmatched deposited-gold intervals contributed IoU=0. Gold-defect region coverage used deposited-gold-derived intervals as the denominator. Residue recall used deposited-gold defect residues as the denominator. Boundary deviation was calculated for matched positive-overlap interval pairs; the median deviation was first computed within each sample-mode and then summarized across eligible sample-modes.

The frozen audit began with 210 sequence-aware sample-modes from the strict audit subset. Six lacked a deposited-gold-derived defect interval, leaving 204 eligible sample-modes for coverage and mean-IoU analysis. Of these, 195 contained at least one positive-overlap interval pair and were eligible for boundary-deviation analysis. Gold-defect region coverage was 93.5%, residue-level recall was 93.0% and mean IoU was 40.9%. The overall median sample-mode boundary deviation was 2.25 residues, displayed as 2.2 residues, with an interquartile range of 0.5–8.375 residues. Means and confidence intervals for the coverage audit were estimated by 5,000 percentile bootstrap resamples at the sample-mode level using seed 20260718.

### Supplementary Methods 3. Staged action registry, legality masks and QC-gated execution

#### Staged execution principle

CryoForge is a staged local-repair system. It does not attempt de novo rebuilding of the complete structure and does not promote a coordinate change solely because an editing program completed. Each candidate passes through region detection, repairability assessment, a legal-action mask, backend-readiness checks, isolated execution and stage-specific post-action QC. Only candidates that satisfy the relevant evidence standard are promoted. Other candidates are left unchanged, rolled back, masked, stopped as data-limited or escalated for expert review.

The stages encode causal dependency rather than simply defect size. Stage0 performs sample-level triage and scope assessment. Stage1 addresses upstream interpretation and connectivity, including short and medium gaps, false links, fragment stitching and conservative overbuilt-like triage. Stage2 addresses backbone geometry, peptide conformation, torsion and local pose only where connectivity is trusted or upstream uncertainty is masked. Stage3 addresses side-chain conformation, local clashes and atom completion after the backbone is stable. Stage4 manages data-limited, unresolved or unsafe regions and provides an explicit manual-review route.

#### Stage1: connectivity and local fragment recovery

Stage1 first applies diagnostic and preflight operations, including boundary-anchor assessment, fragment-graph construction and stitch feasibility. These operations can enable, block or reclassify a repair route but are not counted as coordinate-repair success or failure.

Executable Stage1 action families include conservative false-link breaking, short/medium sequence-aware gap filling, anchored map-first backbone insertion, locally compatible prior-guided low-confidence gap closure, local connectivity regularization and conservative quarantine or removal of clearly unsupported overbuilt-like fragments. Gap filling preserves anchor identity and sequence order. In sequence-aware cases, inserted residues use the corresponding target sequence rather than a generic poly-alanine representation. In sequence-free cases, residue identity can be inferred only when supported by a legal structural or sequence source; otherwise the action is restricted to backbone/connectivity handling or routed to uncertainty management.

Stage1 does not force long-gap reconstruction. Gaps outside the mature local scope, unstable anchors, multiple plausible paths, discontinuous half-map evidence or a prior–map conflict produce a stop, mask or expert-review decision. Connectivity improvement can be accepted even when map fit does not increase, provided that map and half-map support do not regress beyond tolerance, local geometry is safe and neighboring regions remain stable. When density is weak but anchors and a locally compatible prior support one plausible connection, the result can be promoted only as low-confidence partial improvement.

#### Stage2: backbone and local-pose correction

Stage2 operates on regions with adequate upstream connectivity or on independent regions outside unresolved Stage1 masks. Action families include backbone-only real-space refinement, short torsion-window regularization, peptide-flip correction, conservative crankshaft-like backbone adjustment, local rigid-body placement, local-pose real-space refinement and local geometry regularization.

Actions are matched to defect subtype. A peptide flip is not applied to a region dominated by a gap or missing backbone. A local rigid-body action is not used when the principal problem is an unresolved register or chain correspondence. Gap-inserted residues can be regularized in Stage2 when the inserted path is connected but torsion, anchor continuity or local backbone geometry remains suboptimal. Regions without a plausible backbone path are returned to Stage1 or routed to Stage4 instead of being repeatedly refined.

Stage2 post-QC evaluates whether the target backbone defect is reduced or stabilized, whether peptide and torsion geometry remain valid, whether local map/half-map support is non-regressive, and whether neighboring accepted structure is preserved. A candidate that improves geometry while materially reducing local map support is classified as a geometry–map trade-off and is not promoted. If the input region already satisfies the stable-accept threshold, no further edit is required; this avoids treating negligible motion as a failure or repeatedly editing an acceptable model.

#### Stage3: side-chain and local-detail correction

Stage3 receives regions with a stable backbone and a side-chain-level defect. Action families include rotamer correction, clash cleanup, local side-chain real-space refinement, side-chain repacking and completion of missing side-chain atoms. Sequence-aware atom completion uses the known residue identity. When both local map and permitted prior support are weak, a chemistry-based completion action can be considered only to restore chemically required atoms and stereochemistry; it cannot claim map-resolved placement and must be labeled low-confidence.

The Stage3 gate distinguishes completeness from density interpretation. If side-chain atoms are missing, chemically valid completion can be useful even when side-chain density is weak, but the backbone must remain fixed or stable, clashes must not increase and the resulting geometry must be valid. When density is strong, rotamer and placement must additionally be compatible with the map. An attempted completion that cannot identify the intended residue, cannot generate the expected atom set or edits outside the target interval is inapplicable and is excluded before scientific outcome assignment.

#### Stage4: uncertainty management

Stage4 is not a residual failure bin. It records regions for which available evidence does not justify forced automatic repair. Typical causes include weak or discontinuous full- and half-map density, absence of a unique path, unstable gap anchors, overlong gaps, unresolved register or copy correspondence, global/domain placement problems, repeated rollback and backend limitations.

Stage4 outcomes include keep uncertain, retain an unresolved mask, conservatively trim unsupported atoms when independently justified, quarantine an overbuilt-like segment and escalate to expert review. The delivered report records why an action was not attempted, which evidence was missing or conflicting, and whether the region remains a plausible target for manual rebuilding.

#### Legal-action masks and backend readiness

Every action has a registry entry specifying stage eligibility, compatible defect subtypes, required inputs, maximum scope, prior-use policy, risk class, backend, expected output and QC bundle. Before ranking or execution, a deterministic legality mask removes actions that lack required sequence or map inputs, have no available backend, are incompatible with the region type or length, violate a user restriction, overlap an unresolved upstream mask, require an unusable prior, or recently failed in the same region without a changed state.

The same mask is applied to rule-only ranking, GNN-only ranking, GNN/RL ranking and supervisory-language-model review. No decision layer can legalize a masked action. Diagnostic preflight operations are evaluated by their routing value, not by coordinate improvement. An executable action enters the scientific denominator only when it generates an evaluable candidate or a scientifically interpretable stop decision. Pure technical failures and missing-output errors are reported separately.

#### Isolated execution and rollback-safe promotion

Each action is executed in an isolated workspace containing the target model, required local maps or map extracts, legal prior information, the frozen region definition and backend configuration. Outputs are read through the action-specific adapter and verified for coordinate integrity, target contact and expected modification scope before post-QC.

The candidate model is never substituted directly for the current model. Instead, CryoForge compares the candidate with the protected pre-action state. The edit is promoted only if the stage-specific target improves or reaches a stable-accept state without unacceptable regression in map support, half-map evidence, geometry, connectivity or neighboring structure. A failed candidate is discarded and the protected pre-action coordinates remain in the promoted model. Thus, rollback-protected/QC-rejected describes an attempted edit that was not retained; it does not indicate that the delivered model contains the regression.

#### Formal trajectory outcomes and Stage4 escalation

Each released Stage1–Stage3 trajectory receives one of five terminal outcomes:

1. **Standard validated improved:** the stage-specific target improves, all hard safety checks pass and the edit is promoted without a low-confidence qualifier.
2. **Low-confidence partial improvement:** connectivity, completeness or another target improves safely, but weak density, limited prior support or residual uncertainty prevents a high-confidence claim. The promoted interval is labeled for expert awareness.
3. **Data-limited/stopped:** the evidence does not support a unique or safe action; no forced edit is made.
4. **Unchanged/no effective gain:** an action produces no meaningful accepted benefit or the current region already satisfies the stable-accept criterion.
5. **Rollback-protected/QC-rejected:** an attempted edit violates post-QC tolerance or creates an unsafe trade-off and is discarded.

Manual review and expert escalation are Stage4 routing outcomes and are reported with the corresponding Stage4 denominator rather than entered as zero-valued Stage1–Stage3 trajectories. “Final worsened retained” is a safety statistic rather than an additional outcome category. It counts promoted models in which a known harmful candidate survived rollback; no such trajectory was retained in the frozen Fig. 3 outcome corpus.

### Supplementary Methods 4. Learned decision models and supervisory agent

#### Role of the learned decision layer

The learned layer was designed to reduce unnecessary candidate retention and to rank legal repair actions; it did not replace deterministic feasibility or post-action QC. Stage0 used graph-derived sample and region summaries for route prediction. Stage2 and Stage3 used learned actionability and action-ranking models because these stages commonly contain multiple legal local actions with different expected gain and risk. Mature Stage1 gap filling remained predominantly deterministic because anchor validity, gap length, sequence availability and backend readiness impose hard structural constraints that should not be relaxed by a learned score.

#### Graph state representation

For each candidate region, residues were represented as nodes with features describing local map support, Q-score and FSC-Q availability, half-map consistency, backbone and peptide geometry, connectivity, clash/rotamer state, atom completeness, local neighborhood context, unresolved-mask proximity, previous action/QC history and locally permitted prior compatibility. Edges represented sequence adjacency, spatial proximity and relevant local contacts. Region-level and sample-level context included resolution bin, initialization mode, stage, region length, gap length, dominant defect subtype and evidence reliability.

The policy state for region r, action a, stage s and repair round t was represented conceptually as

$$S_{s,t}(r,a) = \{G_r, C_{sample}, C_{stage}, M_a, L_a, H_t\}$$

where G_r is the focal residue graph, C_sample and C_stage are contextual features, M_a is the legal-action mask, L_a is action metadata and H_t is repair history. Deposited-gold features were excluded from runtime states.

#### GNN-only and GNN/RL models

The GNN encoder generated a region embedding used by an actionability head and an action-value head. The GNN-only model was trained by supervised learning to predict candidate actionability, expected positive outcome and relative action suitability from frozen trajectory labels. At inference, it ranked legal actions deterministically from these supervised scores.

The GNN/RL model used the same graph representation and supervised warm start, followed by replay-based value refinement. Replay transitions contained the pre-action state, legal action set, selected action, post-action QC state, terminal outcome, rollback status and reward components. Target-network stabilization and Double-DQN-style action evaluation were used to reduce value overestimation. The learned policy could rank only actions that remained legal after deterministic masking.

The conceptual target value was

$$Q^*(s,a) = R_{quality} + R_{connectivity} + R_{completeness} - P_{risk} - P_{compute} - P_{rollback} - P_{technical}$$

Positive terms rewarded validated stage-specific improvement, safe connectivity recovery, atom completeness and stable map-supported gains. Penalties represented structural risk, unnecessary computation, rollback/QC rejection and technical non-completion. Data-limited stops and preflight-blocked actions were not treated as completed harmful repairs. They supplied actionability and feasibility labels so that the policy could learn when not to select an action.

#### Training, validation and model selection

Training records were derived only from entries in the 600-entry training partition. Validation entries were used for checkpoint selection, probability calibration and operating-point selection. Held-out entries were reserved for final evaluation and were not used to tune rewards, thresholds or action families.

Stage0 checkpoints were selected using route accuracy, macro-F1 and recall for high-risk/global-recovery routes. Stage2 candidate gates were selected under recall constraints to reduce non-improving candidate burden while preserving validated-improved regions. Stage2 and Stage3 action-ranking checkpoints were selected using average precision, oracle-best action match, expected gain and harmful-action rate. Candidate-pool analyses used the frozen development/ablation pools associated with the corresponding source tables; counts of 9,395 Stage2 candidates and 17,219 Stage3 candidates must not be described as held-out repair trajectories unless a separate split-provenance audit establishes that scope.

#### Baselines and ablations

Four decision conditions were compared. Rule-only used the deterministic candidate and default action-priority rules. GNN-only used the supervised graph heads without replay-refined Q values. GNN/RL used the replay-refined contextual action values. Oracle-best was a post hoc upper bound that selected the highest-QC action among evaluated legal actions using outcome information unavailable at runtime.

All conditions used the same candidate pool, legal-action mask, backend-readiness criteria and deterministic post-QC. Therefore, differences between decision models reflect filtering or ranking within the same permitted action space rather than access to different repair tools. Average precision summarized candidate prioritization across thresholds. Matched-recall analyses compared the fraction of non-improving candidates retained at prespecified validated-improvement recall. Oracle-best action match quantified whether the selected legal action agreed with the post hoc highest-utility action. Harmful-action rate measured the fraction of selected executed actions that triggered a true safety regression or rollback category.

#### Supervisory language-model layer

The Codex/GPT-5.5 supervisory layer was a constrained orchestration and reporting component, not a coordinate generator. It received structured region manifests, legal-action masks, backend-readiness status, GNN/RL scores, pre- and post-action QC summaries, rollback history, unresolved masks and user constraints. Its outputs were schema-constrained recommendations such as approve, skip, rollback, accept low confidence, keep uncertain, manual review or recommend an alternative legal action.

The supervisor was invoked when evidence was incomplete or conflicting, predicted gain and predicted risk were both high, an untrained extension action was requested, stage transition required synthesis of multiple unresolved regions, or an auditable expert-facing explanation was required. Routine low-risk actions with complete deterministic evidence did not require language-model arbitration.

Hard limits were enforced outside the language model. The supervisor could not edit PDB/mmCIF coordinates, legalize a masked action, bypass backend-readiness checks, promote a candidate without a validation record, override a hard QC failure, or introduce deposited-gold evidence into runtime decisions. An alternative action recommendation had to select an action already legal and backend-ready. Actions without replay support were labeled rule fallback, user requested or untrained extension rather than being represented as learned-policy decisions.

The supervisory output linked each recommendation to the evidence used, the available alternatives and the governing safety rule. Final reports therefore distinguish accepted edits, low-confidence edits, unchanged regions, rollback-protected candidates, unresolved masks, data-limited regions and expert-review regions. This layer provides traceability and evidence synthesis; coordinate validity remains determined by the registered backend and deterministic QC controller.

### Supplementary Methods 5. Evaluation, statistics and reproducibility

#### Analysis units and denominators

All reported quantities were tied to a predefined analysis unit. An **entry** is one independent cryo-EM map/model case. A **sample-mode** is one entry under one initialization condition. A **candidate region** is a contiguous or locally defined structural target produced by a stage-specific detector. A **stage-entry record** represents a sample-mode or region released into a stage after upstream routing. A **trajectory** is a released stage-specific repair record with an evaluable terminal outcome. An **action record** represents one legal, attempted or preflight-blocked action for a region. A **boundary event** explains why an apparently suspicious region was stopped, assigned low confidence or not promoted.

Counts are not interchangeable across these units. Candidate-region burden is reported per 100 modeled residues:

$$\text{burden} = 100 \times \frac{\text{number of candidate regions}}{\text{number of modeled residues}}$$

Stage-entry candidate counts describe diagnostic burden and need not equal the number of executed repair trajectories. A region can be stopped before execution, and one region can generate more than one legal candidate during exhaustive development. Conversely, a final promoted model can contain accepted edits from several trajectories. Figure captions and source tables therefore state the analysis unit and denominator for each panel.

#### Statistical summaries

Continuous variables were summarized using mean and standard deviation when approximately symmetric and median and interquartile range when skewed. Proportions are accompanied by counts and explicit denominators. Paired analyses were used only when the same entry, sample-mode or region contributed both observations. Bootstrap confidence intervals were resampled at the primary analysis-unit level to avoid treating multiple regions from one sample-mode as independent entries.

For the candidate-region audit, macro summaries were computed across eligible sample-modes. The 95% confidence intervals in the coverage panel were percentile bootstrap intervals from 5,000 sample-mode resamples using seed 20260718. Boundary deviation was summarized by first taking the median among matched interval pairs within each sample-mode and then reporting the distribution of those sample-mode medians. The box represented the interquartile range, the internal line represented the median and whiskers extended to the most extreme values within 1.5 times the interquartile range; points beyond the whiskers were retained as individual observations. No formal p values were used for the primary descriptive repair panels. Where formal related tests are later included, p values will be adjusted within the corresponding metric family using the Benjamini–Hochberg procedure; descriptive counts and source-data summaries remain unadjusted.

#### Frozen repair-trajectory corpus used in Fig. 3

The terminal-outcome analysis supporting Fig. 3 used a frozen pooled full-QC repair corpus containing 26,153 released Stage1–Stage3 trajectories. This corpus was assembled for action-registry maturation and descriptive QC analysis across the benchmark workflow. It is distinct from the 129-entry held-out test partition and is not used as a held-out generalization estimate. The corresponding source table retains the source partition, sample-mode, stage, target region, selected action, post-QC decision and rollback state needed to audit each trajectory.

The five terminal outcomes were: 22,191 standard validated improvements (84.9%), 970 low-confidence partial improvements (3.7%), 2,192 data-limited/stopped trajectories (8.4%), 689 rollback-protected/QC-rejected trajectories (2.6%) and 111 unchanged/no-effective-gain trajectories (0.4%). Standard and low-confidence positive outcomes together accounted for 23,161 trajectories (88.6%). No final worsened trajectory was retained. Stage4/manual-review routing is summarized separately and is not represented by a zero-valued row in this Stage1–Stage3 trajectory denominator.

The stage-specific local-gain panels use their own frozen analysis units. Stage1 connectivity was evaluated across 5,443 target gaps; unresolved connectivity breaks decreased from 4.22 to 0.08 per target gap, corresponding to a 98.1% reduction. Stage2 geometry was evaluated across 1,127 repair regions; geometry defects decreased from 1.38 to 0.16 per region, corresponding to an 88.4% reduction. Stage3 local-detail performance was evaluated across 53,672 evaluable edits, of which 70.8% achieved improved or partially improved status. These denominators are stage-specific and are not additive.

#### Local repair metrics

Stage1 connectivity was evaluated using break burden before and after the accepted repair, anchor continuity, gap closure, inserted-residue count and unresolved-gap status. Stage2 used geometry defects per region, peptide and torsion state, target displacement, map/half-map non-regression and neighbor stability. Stage3 used side-chain completeness, rotamer and clash state, local density compatibility when informative and backbone stability.

Global aligned Cα RMSD to the deposited gold model was a post hoc scaffold-preservation measure. In the frozen paired subset used in Fig. 3, mean aligned Cα RMSD was 0.64 Å before repair and 0.63 Å after post-QC promotion ($n=13$ paired models; mean $\Delta=-0.008$ Å). It was calculated after compatible chain or local correspondence and did not participate in runtime repair. Because CryoForge performs local repair rather than global rebuilding, a small global RMSD change can coexist with substantial local connectivity, geometry or completeness improvement. Conversely, an apparent RMSD decrease cannot rescue an action that fails map, geometry or safety QC.

#### External-initialization evaluation

Classic and CryoAtom-derived initial models were compared on the same 129 held-out entries whenever both outputs and coordinate-compatible maps were available. Entry-level metrics included Cα breaks per 1,000 modeled Cα atoms, deposited-gold Cα coverage, matched modeled Cα fraction, unmatched modeled Cα fraction and compatible aligned RMSD. Stage-entry phenotype counts were analyzed separately from repair trajectories. Large candidate pools can be reduced sharply by repairability, legality and post-QC gates; therefore, diagnostic candidate counts and promoted Stage2/Stage3 trajectories are reported as distinct quantities.

CryoAtom analyses were interpreted as an external-initialization compatibility test. They support the conclusion that the same evidence-gated framework can triage and conservatively process a different initialization phenotype. They do not establish universal generalization to all initial-model generators.

#### Seeds, checkpoints and software records

Random seeds were recorded for benchmark splitting, graph-model initialization, minibatch order, replay sampling, action-ranking tie breaks, bootstrap resampling and stochastic backends when a seed was exposed. Deterministic operations were marked deterministic rather than assigned an artificial seed. If an external program used hidden stochastic behavior, this limitation was recorded in the environment manifest.

Checkpoints were frozen before held-out evaluation. Each record contains the stage, feature-schema version, training and validation partitions, seed, selected epoch or step, validation criterion and checksum. Stage0 models were selected using route accuracy, macro-F1 and high-risk-route recall. Stage2 and Stage3 models were selected using recall-preserving candidate reduction, average precision, oracle-action agreement and harmful-action rate. Held-out labels were not used for checkpoint or threshold selection.

Run manifests record the CryoForge revision, action-registry and QC-policy versions, Python environment, machine-learning libraries, map and structure-processing libraries and external backends. The primary initialization versions were ModelAngelo v1.0.16 (commit ffb872cb) and PHENIX 2.0-5936-cuda12. Exact versions of Coot, Servalcat/REFMAC, MolProbity-related validation, Q-score, FSC-Q and auxiliary packages are provided in the frozen software manifest distributed with the reproducibility package.

#### Provenance and reproducibility

For each major experiment, CryoForge recorded the dataset and split manifests, region and action manifests, QC summaries, repair trajectories, replay-buffer version, checkpoint manifest, action-registry version, QC-policy version, seed record, software manifest and figure source tables. Internal filesystem paths, shard logs and raw prompts are not required to interpret the method and are not included in the narrative Supplementary Information. They remain internal provenance records where permitted.

The public release provides code, configuration templates, schema descriptions, sample-download utilities and representative report examples without redistributing original cryo-EM maps or deposited coordinate files. Runtime reports identify accepted, low-confidence, unchanged, rolled-back, stopped and expert-review regions and preserve the evidence used for each decision. Deposited-gold annotations are exposed only in post hoc audit outputs.

### Supplementary Figure captions

#### Supplementary Figure 1. Evidence hierarchy and stage routing in CryoForge

Residue-level map, half-map, geometry, connectivity, completeness and neighborhood evidence is converted into candidate regions. Detection is followed by a separate repairability gate that assigns map-supported repair, locally compatible prior-guided low-confidence repair, ambiguity, data limitation or manual review. Repairable regions enter Stage1 connectivity recovery, Stage2 backbone/local-pose correction or Stage3 side-chain/local-detail correction according to upstream dependency. Data-limited or unsafe regions can enter Stage4 from any stage. Deposited-gold information is excluded from runtime routing and shown only in the separate post hoc audit branch.

#### Supplementary Figure 2. Runtime evidence boundary and provenance-verified outcome accounting

Runtime actions use full-map and half-map evidence, local QC, geometry, connectivity, legal sequence/prior information, action history and learned scores. Candidate coordinates are generated in isolated workspaces and are promoted only after deterministic stage-specific QC; rejected candidates are rolled back. The pooled full-QC trajectory corpus used in Fig. 3 is retained with source-partition and sample-mode provenance and is interpreted as a descriptive action/QC analysis. Independent held-out analyses use the frozen 129-entry test partition. No deposited-gold evidence entered runtime decisions.

### Supplementary Tables

#### Supplementary Table 1. Benchmark design and split structure

| Item | Definition | Count |
|---|---|---|
| Independent benchmark unit | Cryo-EM entry | 849 |
| Resolution 2.5–3.0 Å | Entry-level stratum | 283 |
| Resolution 3.0–3.5 Å | Entry-level stratum | 283 |
| Resolution 3.5–4.0 Å | Entry-level stratum | 283 |
| Training partition | Frozen entry-level split | 600 |
| Validation partition | Frozen entry-level split | 120 |
| Held-out test partition | Frozen entry-level split | 129 |
| Sequence-aware sample-modes | One per entry | 849 |
| Sequence-free sample-modes | One per entry | 849 |
| Total sample-modes | Both initialization modes | 1,698 |
| Held-out external-initialization cohort | Frozen test entries used for CryoAtom compatibility analysis | 129 |

#### Supplementary Table 2. Frozen evidence and routing thresholds

| Evidence or rule | Frozen criterion | Interpretation |
|---|---|---|
| Supported residue Q-score | Q-score ≥0.40 | Residue-level density-support indicator |
| Low-density FSC-Q trigger | Mean FSC-Q <0.25 or minimum FSC-Q <0.10 | Routes region to weak-density review when reliable |
| FSC-Q discrepancy-support flag | \|FSC-Q\| ≤0.50 | Policy-specific discrepancy criterion |
| Hard peptide break | C–N >2.00 Å | Stage1 connectivity signal |
| Expected peptide C–N | 1.15–1.55 Å | Local connectivity reference interval |
| Hard Cα break | Adjacent Cα–Cα >5.00 Å | Stage1 connectivity signal |
| Expected adjacent Cα | 3.40–4.20 Å | Local backbone interval |
| Suspicious peptide ω | 35° < \|ω\| <145° | Peptide-geometry signal |
| Severe non-neighbor clash | Heavy-atom distance <1.60 Å | Geometry/clash signal |
| Candidate merge | Same chain; selected sequence separation ≤2 | Region construction |
| Routine map-first gap fill | 1–8 missing residues | Mature high-confidence scope |
| Conditional prior-guided gap fill | 9–12 missing residues | Low-confidence/review-dependent scope |

| Evidence or rule | Frozen criterion | Interpretation |
| --- | --- | --- |
| Overlong local gap | >12 missing residues | Stage4/global/expert route |
| Local prior support | ≥3 Cα pairs and coverage ≥0.30 | Minimum local prior compatibility |
| Strict Stage0 prior support | Adaptation passed and coverage ≥0.45 or ≥4 Cα pairs | Prior eligible for stronger weighting |
| Post hoc defect seed | Nearest deposited-gold Cα distance ≥3.0 Å | Offline audit only |
| Post hoc audit eligibility | ≥50 mappable Cα atoms | Offline audit only |

#### Supplementary Table 3. Stage-specific action families and promotion criteria

| Stage | Primary target | Representative action families | Minimum release condition | Promotion criterion |
| --- | --- | --- | --- | --- |
| Stage0 | Scope and route | Triage, coordinate-frame check, global/scope review | Valid map/model workspace | Route assigned without deposited-gold evidence |
| Stage1 | Connectivity, gaps, local fragments | False-link break, anchored gap fill, sequence-aware fill, map-first insertion, locally compatible prior-guided fill, conservative overbuilt-like quarantine | Stable anchors or interpretable connectivity target; legal backend | Connectivity/gap target improved; geometry safe; map/half-map non-regressive; neighbor stable |
| Stage2 | Backbone, peptide, torsion, local pose | Backbone-only RSR, torsion regularization, peptide flip, local rigid-body fit, local-pose RSR | Connectivity trusted or unresolved interval masked | Backbone/geometry target improved or stable-accepted; no unsafe map or neighbor regression |
| Stage3 | Side chain, clash, atom completeness | Rotamer correction, clash cleanup, side-chain RSR/repacking, atom completion | Backbone stable; residue identity and target resolvable | Completeness/rotamer/clash improved; backbone stable; map-compatible when informative |
| Stage4 | Data limitation and unresolved risk | Keep uncertain, unresolved mask, quarantine, conservative trim, manual review | Repairability gate blocks safe automated repair | No forced promotion; evidence and review reason recorded |

#### Supplementary Table 4. Frozen pooled full-QC repair outcomes used in Fig. 3

| Outcome | n | Percentage |
| --- | ---: | ---: |
| Standard validated improved | 22,191 | 84.9% |
| Low-confidence partial improvement | 970 | 3.7% |
| Data-limited / stopped | 2,192 | 8.4% |
| Rollback-protected / QC-rejected | 689 | 2.6% |

| Outcome | n | Percentage |
|---|---|---|
| Unchanged / no effective gain | 111 | 0.4% |
| Total released trajectories | 26,153 | 100.0% |

The analysis unit is a released Stage1–Stage3 repair trajectory from the pooled full-QC action/QC corpus. It is not an entry-level or held-out-test denominator. Standard and low-confidence positive outcomes sum to 23,161 trajectories (88.6%). Stage4/manual-review routes are reported separately. No final worsened trajectory was retained.

#### Supplementary Table 5. Analysis units and principal evaluation metrics

| Analysis unit | Typical question | Principal metrics | Required denominator statement |
|---|---|---|---|
| Entry | How many independent cryo-EM cases were studied? | Split count, resolution distribution, paired builder comparison | Number of unique entries |
| Sample-mode | How does initialization condition affect burden or routing? | Candidate burden, Stage0 route, QC availability | Entries × initialization mode |
| Candidate region | What defect types were detected and judged repairable? | Defect composition, repairability, unresolved masks | Number of stage-specific regions |
| Stage-entry | What reached a stage after upstream gates? | Stage-entry burden and release rate | Eligible sample-modes or regions at that stage |
| Trajectory | What was the terminal consequence of a released repair path? | Validated, partial, stopped, unchanged, rollback, manual review | Released evaluable trajectories |
| Action | Did the policy select a useful legal action? | Oracle match, harmful-action rate, action regret | Legal/evaluated action records |
| Boundary event | Why was an apparent target not promoted? | Weak density, long gap, trade-off, no legal action, ambiguity | Stage-specific non-improving or low-confidence events |
| Post hoc deposited-gold interval | Did candidate proposals cover benchmark defects? | Region coverage, residue recall, IoU, boundary deviation | Eligible audit sample-modes or gold-derived intervals |
